# Protein kinase Cδ is essential for the IgG response against T cell-independent type 2 antigens and commensal bacteria

**DOI:** 10.1101/2021.07.20.453051

**Authors:** Saori Fukao, Kei Haniuda, Hiromasa Tamaki, Daisuke Kitamura

## Abstract

Antigens (Ags) with multivalent and repetitive structure elicit IgG production in a T cell-independent manner. However, the mechanisms by which such T cell-independent type-2 (TI-2) Ags induce IgG responses remain obscure. Here we report that BCR engagement with a TI-2 Ag but not with a T cell-dependent (TD) Ag was able to induce the transcription of *activation-induced cytidine deaminase* (*AID*) and efficient class switching to IgG3 upon co-stimulation with IL-1 or IFN-α. TI-2 Ags strongly induced the phosphorylation of protein kinase C (PKC)δ and PKCδ mediated the *AID* transcription through the induction of BATF, the key transcriptional regulator of AID expression. In PKCδ-deficient mice, production of IgG was intact against TD Ag but abrogated against typical TI-2 Ags as well as commensal bacteria, and experimental disruption of the gut epithelial barrier resulted in fatal bacteremia. Thus, our results revealed novel molecular requirements for class-switching in the TI-2 response and highlighted its importance in homeostatic commensal-specific IgG production.

## Introduction

Ag-specific antibody production is essential for humoral immunity. After TD Ag exposure, B cells are activated by interacting with cognate T cells and then proliferate, undergo class switching, and differentiate into plasma cells (PCs) and memory cells. In contrast, TI Ags activate B cells without cognate help of T cells. TI type 1 (TI-1) Ags engage Toll-like receptors (TLRs) in addition to the B cell receptor (BCR) whereas TI-2 Ags extensively crosslink the BCR because of their highly repetitive structure (Mond et al., 1995). In addition, some help from non-T cells, such as dendritic cells and innate lymphoid cells, may support B cells in a TI response (Magri et al., 2014; Balázs et al., 2002). Following activation, TI Ags induce proliferation, class-switching and antibody production by B cells. TI-2 Ags are large multivalent molecules, such as bacterial capsular polysaccharides and viral capsids, and thus antibody production against polysaccharides by the TI-2 response confers protection against disease (Mond et al., 1995; Lesinski and Julie Westerink, 2001). To date, however, the mechanism of B cell activation in the TI-2 response is poorly understood as compared to that in TD and TI-1 responses.

Engagement of the BCR by Ag activates various signaling cascades and promotes B cell survival, proliferation and differentiation. The antibody production in a TI-2 response, but not in a TD response, depends upon proximal BCR signaling molecules such as Btk and BLNK (Khan et al., 1995; Xu et al., 2000; Fruman et al., 2000), suggesting that BCR signaling plays a more critical role for B cell activation in a TI-2 response than in a TD response. Given the highly repetitive structure of TI-2 Ags and the high demands for BCR signaling in TI-2 responses, TI-2 Ags seem to induce BCR signaling and the subsequent response more strongly than TD Ags. However, such functional differences between TD Ags and TI-2 Ags have not been investigated.

Class (isotype) switching of the immunoglobulin (Ig) on B cells from IgM to either IgG, IgE, or IgA is caused by selective recombination of Igh constant region (C_H_) genes, namely, class-switch recombination (CSR). Selection of the C_H_ gene to be recombined is determined by T cell-derived cytokines in the TD response but the mechanism in the TI-2 response is not clear. Immunization with TI-2 Ags, including NP-Ficoll and bacterial polysaccharides, predominantly elicits IgG3 (Perlmutter et al., 1978; Slack et al., 1980; Rubinstein and Stein, 1988) and IgG3 is required for protection against pneumococcal infection (McLay et al., 2002). CSR absolutely requires AID (Muramatsu et al., 2000). AID deaminates deoxycytidine and introduces DNA double strand breaks (DSB) in the switch (S) regions lying 5’ of the Cμ gene and the other targeted C_H_ gene by triggering the DNA repair machinery (Stavnezer et al., 2008). CSR proceeds through looping-out deletion of the DNA segment intervening between Sμ and the other S region from the chromosome and re-ligation of the DSB free ends in the two S regions. Consequently, it leads to replacement of the Cμ gene with a different C_H_ gene downstream of the variable region exon in the *Igh* locus.

In a TD immune response, stimulation with CD40L and cytokines, such as IL-4 and TGFβ, induce the expression of AID and subsequent CSR in the B cell (Vaidyanathan et al., 2014; Dedeoglu, 2004), while signaling through TLRs, BCR and TACI induce the expression of AID and CSR in a TI-1 response (Xu et al., 2012). On the other hand, the signaling and molecular demands for the induction of AID and CSR in the TI-2 response are far less understood, presumably due to the lack of *in vitro* studies that mimic a TI-2 response. Although TACI is required for IgG production in a TI-2 response (von Bülow et al., 2001), stimulation of TACI alone induces the expression of AID and IgG production only modestly (Castigli et al., 2005). Therefore, signaling through other receptors besides TACI seems to be required. As antibody production itself is disrupted in mice lacking proximal BCR signaling molecules (Khan et al., 1995; Xu et al., 2000), the requirement of BCR signaling and the downstream molecules for CSR remain unclear. Here, we report our finding that stimulation of the BCR with NP-Ficoll, a typical TI-2 Ag, triggered IgG CSR in the presence of secondary stimulation by IL-1, IFNα or TLR ligands *in vitro* and that PKCδ is critical for the Ag-mediated upregulation of AID and IgG production in the TI-2 response.

## Results

### A TI-2 Ag induces B-cell proliferation and potentiates CSR to IgG

Considering the unique structure of TI-2 Ags, it is plausible that the engagement of the BCR with TI-2 Ags and TD Ags differently induces downstream signaling that leads to B cell activation, although this idea has not been tested properly so far. We tested this *in vitro* by stimulating NP-specific B cells with the TD Ag NP-CGG or the TI-2 Ag NP-Ficoll to compare their ability to induce signaling, proliferation and antibody production. NP-specific B cells were prepared from *Igκ ^-/-^* mice (expressing only λ isotype light chain) carrying a *V_H_ B1-8* knock-in gene encoding a V_H_ region which binds to NP when coupled with a λ light chain. Although NP-CGG induced little proliferation and no IgM production, NP-Ficoll induced strong proliferation and IgM production, whereas neither induced IgG production (Figure 1, A and B). Thus, although NP-Ficoll alone can strongly activate B cells, additional stimulation seemed to be required for induction of class switching, similarly to a previous report that anti-δ mAb/dextran and TLR ligands synergistically induced AID and CSR (Pone et al., 2012). Indeed, NP-Ficoll induced IgG production, generation of IgG^+^ cells and *Aicda* transcription in the presence of TLR ligands, LPS, R-848 or CpG, while NP-CGG did so marginally (Figure 1 - figure supplement 1, A-C). Besides TI-1 Ags, such as TLR ligands, we sought to identify co-stimulating molecules that induce class switching in the TI-2 response, some of which were suggested previously (Magri et al., 2014; Balázs et al., 2002). Since IgG3 is the most dominant class-switched Ig isotype produced in a TI-2 response, we screened various cytokines for their ability to promote the production of IgG3 in the presence of NP-Ficoll, and identified IL-1α, IL-1β, and IFNα as efficient co-stimuli for IgG3 production (Figure 1 - figure supplement 1D and Figure 1 C). These cytokines, together with NP-Ficoll, induced generation of IgG3^+^ cells (Figure 1 D), as well as IgG1^+^ and IgG2b^+^ cells to a lesser extent (Figure 1 - figure supplement 1E). In the presence of these cytokines, NP-Ficoll was far more potent than NP-CGG for IgG3 production, IgG3^+^ cell generation, Sµ-Sγ3 CSR (detectable as the Iγ3-Cµ transcript from the switch circle DNA), and the induction of *Aicda* transcription (Figure 1, C-E). NP-Ficoll or each of these cytokines alone could not induce IgG3 and *Aicda* transcription (data not shown, Figure 1 - figure supplement 1F) (Xu et al., 2012; Kinoshita et al., 2001). Collectively, these results indicate that BCR signaling elicited by TI-2 Ag-engagement is pivotal for the TI-2 B-cell response, namely induction of proliferation and antibody production, as well as potentiation of CSR to IgG.

**Figure 1.**
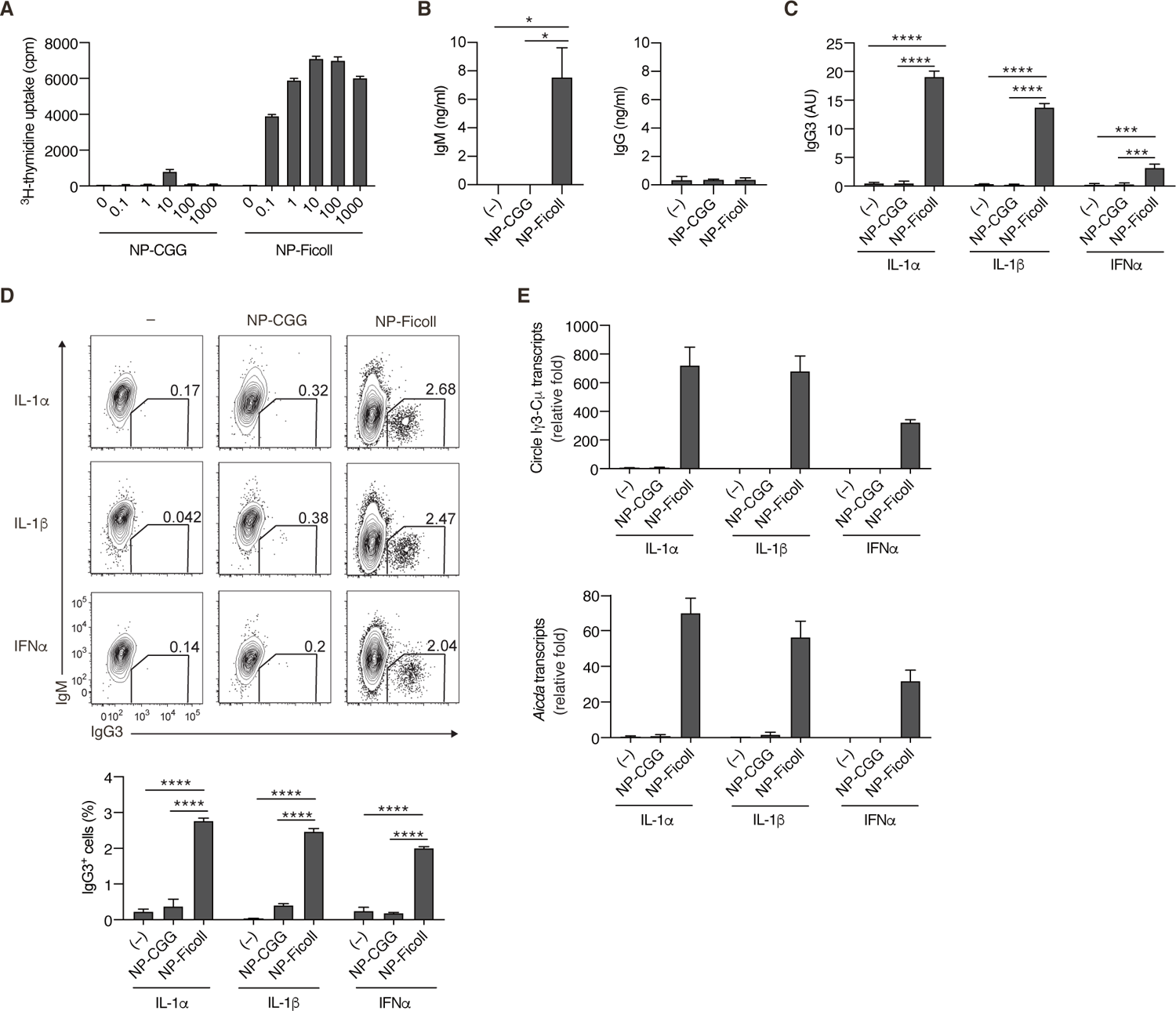
TI-2 Ag distinctively induces B cell activation and CSR to IgG *in vitro*. **(A and B)** Naïve B cells purified from the spleen of *Igκ^-/-^ B1-8^flox/+^* mice were stimulated with NP-CGG or NP-Ficoll. (A) ^3^H-thymidine incorporation on day 3. (B) IgM and IgG concentrations in the culture supernatants on day 3. **(C-E)** *Igκ^-/-^ B1-8^flox/+^* B cells were stimulated with none (-), NP-CGG or NP-Ficoll and IL-1α, IL-1β, or IFNα. (C) ELISA of IgG3 in the culture supernatant on day 3. AU, arbitrary units. (D) Representative flow cytometric plots on day 3 with the numbers indicating percentages of IgG3^+^ B cells (top). The frequencies of IgG3^+^ cells among the B cells (bottom). (E) qRT-PCR analysis of the circle Iγ3-Cµ and *Aicda* transcripts on day 2. Data are means ± SDs of 2 (B, C), 2-3 (D) or 3 (A) biological replicates or 3 technical replicates (E). The data are representative of at least 3 (A and B) or 2 (C-E) independent experiments. *, p<0.05; ***, p<0.001; ****, p<0.0001; p values were calculated by 1 (B) or 2 (C, D)-way ANOVA with Tukey’s test. The following source data and figure supplement are available for Figure 1: Source data 1. Source data for Figure 1A-E. Figure supplement 1. Supplementary data for Figure 1.

### PKCδ is required for IgG production and AID expression induced by TI-2 Ag stimulation

By Western blot analyses for BCR signaling using NP-specific B cells, we noticed that NP-Ficoll induced phosphorylation of PKCδ tyrosine 311 (Y311), indicative of activation of the kinase (Balasubramanian et al., 2006), more strongly than NP-CGG (Figure 2 A). PKCδ belongs to a novel PKC subfamily (Salzer et al., 2016), whose function in the immune response remains elusive. Thus, we analyzed the role of PKCδ in the TI-2 response using PKCδ-deficient (*Prkcd^-/-^*) mice. B cells from *Prkcd^-/-^* mice normally proliferate and produce IgM when stimulated with NP-Ficoll alone (Figure 2 - figure supplement 1, A and B). Upon co-stimulation with NP-Ficoll and IL-1α, IL-1β, and IFNα, *Prkcd^-/-^* B cells produced comparable levels of IgM but markedly reduced levels of IgG3 compared to *Prkcd^+/+^* cells (Figure 2 B). As class switching occurs along with cell division (Deenick et al., 1999), we analyzed the frequency of IgG3^+^ cells at each cell division using B cells labelled with CellTrace violet (CTV). While cell division was almost equivalent between these cells after co-stimulation with any of the cytokines and NP-Ficoll (Figure 2 - figure supplement 1C), the frequencies of IgG3^+^ cells barely increased at any points of cell division in *Prkcd*^-/-^ cells in contrast to *Prkcd*^+/+^ cells (Figure 2 C). Therefore, PKCδ is necessary for generation of IgG3^+^ cells regardless of cell division. Accordingly, *Igh* locus CSR to IgG3 assessed by the circle Iγ3-Cµ transcript and the expression of *Aicda* transcripts were attenuated in *Prkcd^-/-^* B cells (Figure 2 D). Collectively, these results suggest that PKCδ mediates the expression of AID and class switching to IgG3 induced by BCR-stimulation with TI-2 Ag and IL-1/IFNα co-stimulation.

**Figure 2.**
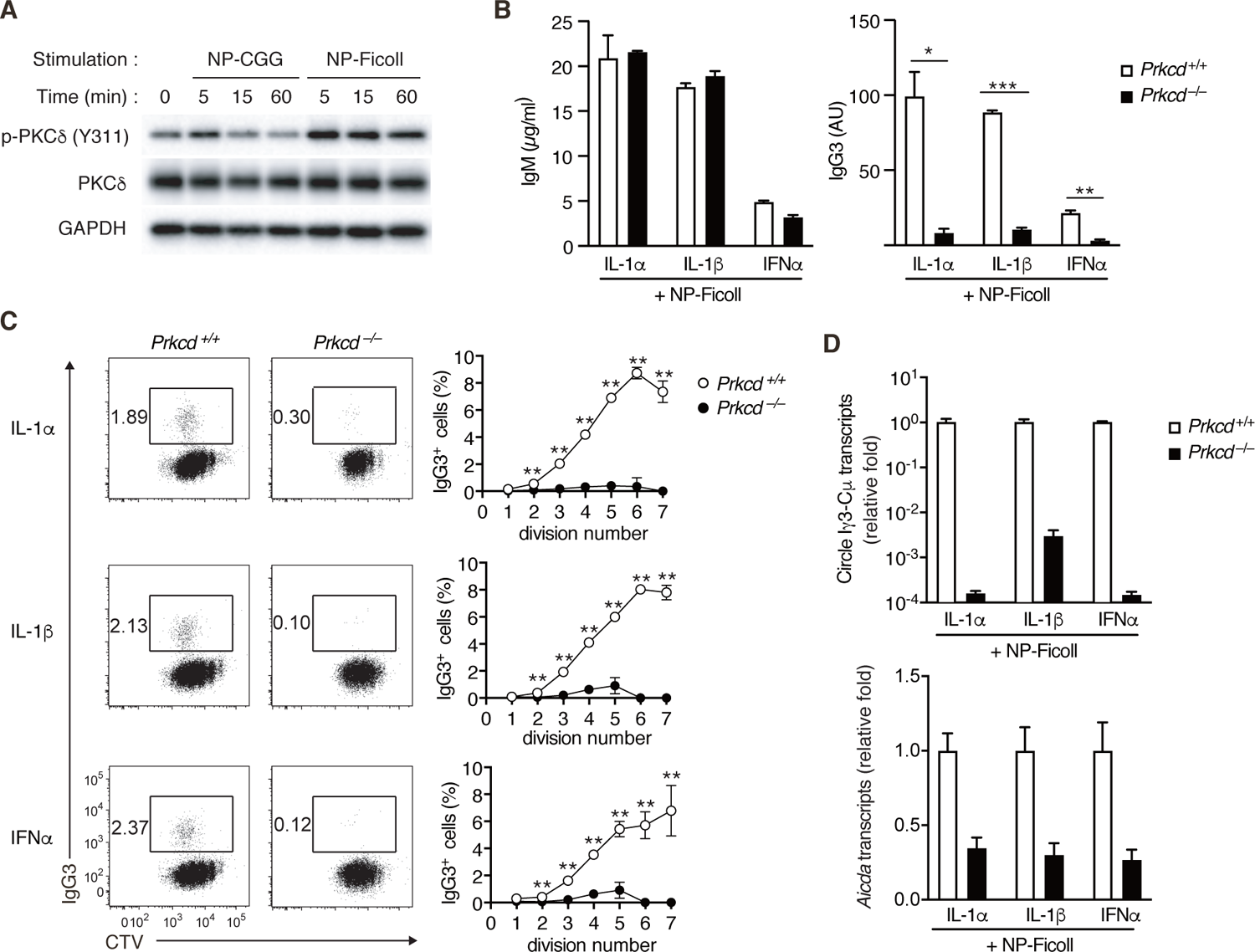
Impaired TI-2 Ag-mediated CSR in PKCδ-deficient B cells. **(A)** Immunoblot analysis of the indicated molecules in *Igκ^-/-^ B1-8^flox/+^* B cells stimulated with NP-CGG or NP-Ficoll for the indicated time periods. **(B-D)** *Prkcd* ^+*/+*^ *Igκ^-/-^ B1-8^flox/+^* or *Prkcd ^-/-^ Igκ^-/-^ B1-8^flox/+^* B cells were stimulated with NP-Ficoll and IL-1α, IL-1β, or IFNα. The B cells were labeled with CTV before culture in C. (B) ELISA of IgM and IgG3 in the culture supernatants on day 5. AU, arbitrary units. (C) Representative flow cytometric plots of the B cells on day 3 showing percentages of IgG3^+^ B cells (left). The frequencies of IgG3^+^ cells at each cell division number (right). (D) qRT-PCR analysis of the circle Iγ3-Cµ and *Aicda* transcripts on day 2. Data are means ± SDs of 2 (B) or 3 (C) biological replicates or 3 technical replicates (D). The data are representative of at least 3 (A and B) or 2 (C and D) independent experiments. *, p<0.05; **, p<0.01; ***, p<0.001; p values were calculated by unpaired Multiple t test (B, C). The following source data and figure supplement are available for Figure 2: Source data 1. Source data for Figure 2B-D; Source data 2-5. Source data for Figure 2A. Figure supplement 1. Supplementary data for Figure 2.

*Prkcd^-/-^* B cells also exhibited defects in the production of IgG and *Aicda* transcripts, but normal IgM production, upon stimulation with NP-Ficoll and TLR ligands (Figure 2 - figure supplement 1, D and E), but no defects in the production of IgM, IgG, and *Aicda* transcripts, upon stimulation with TLR ligands alone (Figure 2 - figure supplement 1, F and G). Together with the above results, PKCδ appears to be selectively required for TI-2-Ag-mediated BCR signaling to potentiate class switch recombination to produce IgG.

### PKCδ mediates IgG production in a TI-2 response *in vivo*

We next evaluated the contribution of PKCδ to IgG production in a TI-2 immune response using mice carrying loxP-flanked *Prkcd* alleles and *Cd19*-cre allele that lack *Prkcd* specifically in B cells (referred to herein as *Cd19^cre/+^ Prkcd^f/f^*). We first examined the cellularity of mature B cell subpopulations in such mice and their control (*Cd19^cre/+^ Prkcd^+/+^*). The numbers of follicular and marginal zone B cells in spleens were comparable between these mice, whereas those of splenic and peritoneal B1 cell populations were slightly increased in *Cd19^cre/+^ Prkcd^f/f^* mice (Figure 3 - figure supplement 1A). Thus, PKCδ is dispensable for the development of B cells, as previously described (Mecklenbräuker et al., 2002; Miyamoto et al., 2002) To analyze the role of PKCδ in a TI-2 response, we immunized *Cd19^cre/+^ Prkcd^f/f^* and the control mice with NP-Ficoll. Although the production of serum anti-NP IgM was slightly enhanced in *Cd19^cre/+^ Prkcd^f/f^* mice, that of anti-NP IgG3 was severely suppressed (Figure 3 A), and so was that of anti-NP IgG1, IgG2b, and IgG2c (Figure B). We then analyzed PCs in the spleen one week after immunization. Among the NP-specific PCs, the proportion and the number of IgG3- and IgG2b-producing PCs were severely decreased in *Cd19^cre/+^ Prkcd^f/f^* mice compared to the control mice, whereas the number of IgM^+^ PCs was comparable between the two groups (Figure 3 C). Therefore, PKCδ is required for the generation of IgG PCs and the subsequent production of IgG in the TI-2 response.

**Figure 3.**
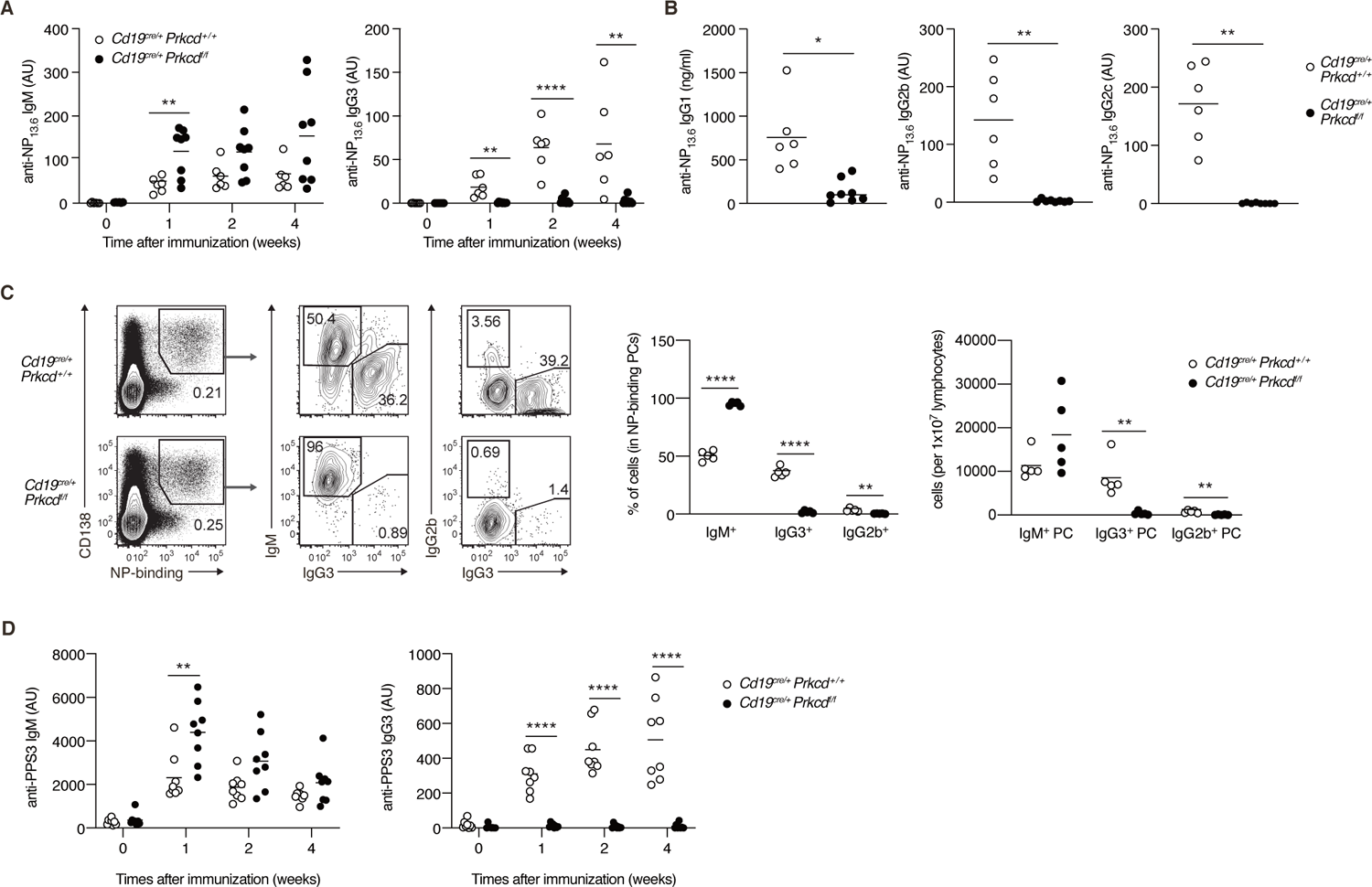
PKCδ is required for IgG production in TI-2 response *in vivo*. **(A-C)** *Cd19^cre/+^ Prkcd^+/+^* and *Cd19^cre/+^ Prkcd ^f/f^* mice were immunized with NP-Ficoll. (A) ELISA of serum anti-NP IgM and IgG3 at the indicated weeks. (B) ELISA of serum anti-NP IgG1, IgG2b and IgG2c at 2 weeks after immunization. (C) Representative flow cytometric plots of the spleen cells on day 7 after immunization. The numbers indicate the percentage of cells in each gate (left). The frequencies of IgM^+^, IgG2b^+^ and IgG3^+^ cells among NP-binding PCs (NP^+^ CD138^+^) (middle) and the numbers of such cells per 1×10^7^ total lymphocytes (right) are plotted. **(D)** ELISA of anti-PPS3 IgM and IgG3 in the serum of *Cd19^cre/+^ Prkcd^+/+^* and *Cd19^cre/+^ Prkcd ^f/f^* mice at the indicated weeks after immunization with PPS3. Results are presented in AU, arbitrary units (A, B, D). Small horizontal bars are the means of 6-8 (A amd B), 5 (C) and 8 (D) biological replicates. Each symbol represents an individual mouse. The data are representative of 3 (A and C) or 2 (B and D) independent experiments. **, p<0.01; ***, p<0.001; ****, p<0.0001; p values were calculated by unpaired Multiple (A, C, D) or 2-tailed unpaired Welch’s t test (B). The following source data and figure supplement are available for Figure 3: Source data 1. Source data for Figure 3A-D. Figure supplement 1. Supplementary data for Figure 3.

The capsular polysaccharides of S. pneumoniae, such as pneumococcal polysaccharide serotype 3 (PPS3), are also classified as TI-2 Ags, and immunization with PPS3 is known to induce an Ag-specific IgG3 response (McLay et al., 2002). Thus, we next immunized *Cd19^cre/+^ Prkcd^f/f^* and control mice with PPS3. Although early production of anti-PPS3 IgM was modestly enhanced in *Cd19^cre/+^ Prkcd^f/f^* mice, production of anti-PPS3 IgG3 was ablated in these mice (Figure 3 D). Thus, PKCδ appears to be generally required for IgG production in response to a variety of TI-2 Ags.

We next assessed whether PKCδ is also required for IgG production in a TD response. After immunization with NP-CGG, the production of anti-NP IgM was transiently enhanced in *Cd19^cre/+^ Prkcd^f/f^* mice at one week, whereas anti-NP IgG1 and IgG3 titers were normal in these mice (Figure 3 - figure supplement 1B). These data indicated that PKCδ signaling is required for IgG production in a TI-2 response, but not in a TD response.

### PKCδ mediates class switching through induction of AID in a TI-2 response *in vivo*

We next assessed whether PKCδ mediates IgG production through class switching in an *in vivo* TI-2 response. To discriminate Ag-specific B cell responses, we transferred CTV-labelled naïve B cells from *B1-8^hi^* CD45.1 mice into C57BL/6 (B6) mice, which were immunized with NP-Ficoll on the next day and analyzed by flow cytometry 3 days later. About 30% of the splenic donor cells were IgG3^+^ in the mice transferred with control B cells, whereas only about 3% were IgG3^+^ in recipients of *Cd19^cre/+^ Prkcd^f/f^* B cells (Figure 4 A). IgG3^+^ cells emerged at the second cell division and their frequency was increased as the cell division proceeded in the cells of control mice, whereas the frequency of IgG3^+^ cells was extremely low in the cells of *Cd19^cre/+^ Prkcd^f/f^* mice at any point of cell division (Figure 4 A). Thus, PKCδ mediates the generation of IgG3^+^ cells in a manner unrelated to cell division.

**Figure 4.**
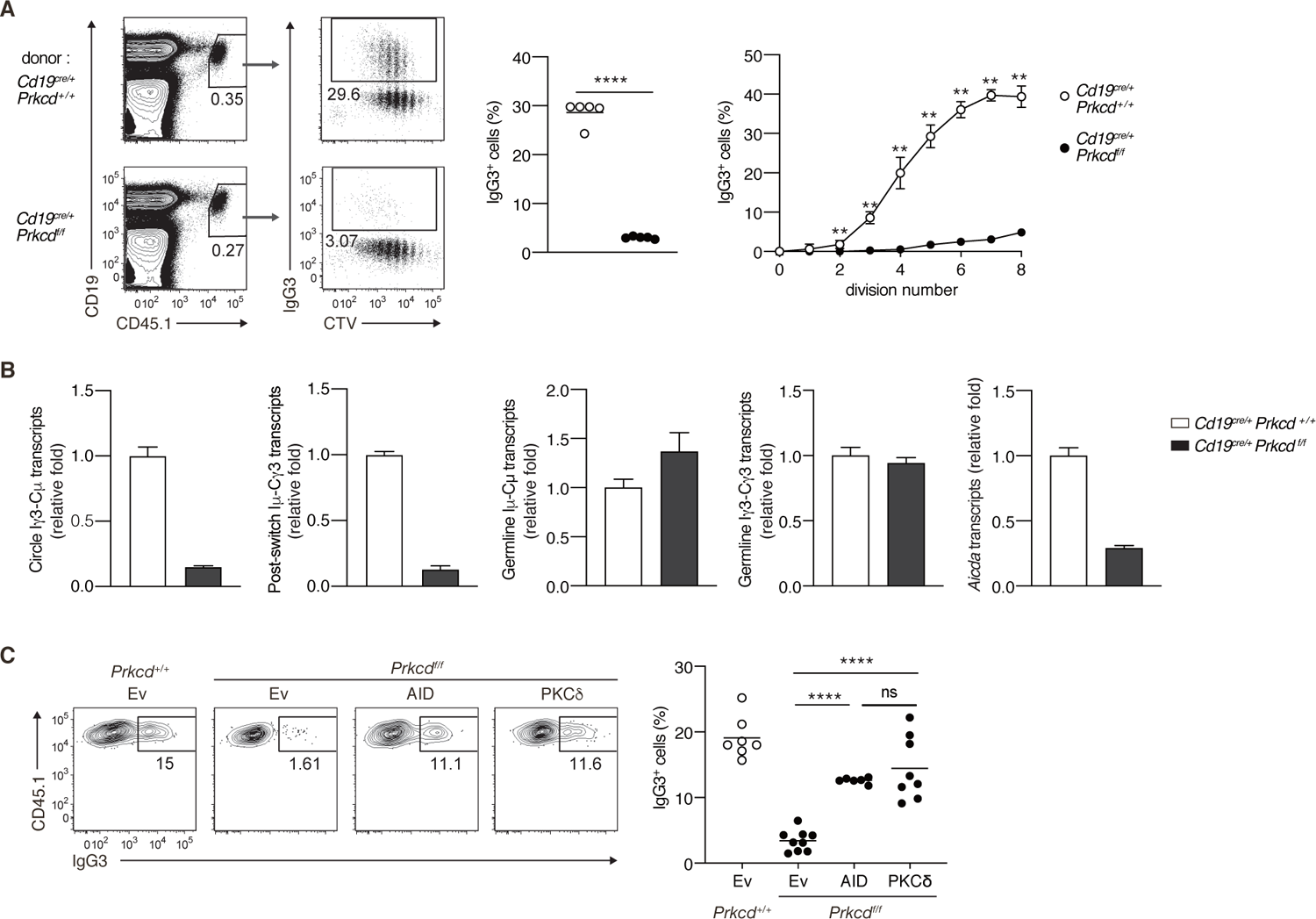
PKCδ signaling promotes IgG class switching by inducing AID. **(A and B)** B cells purified from *Prkcd^+/+^ Cd19^cre/+^ B1-8^hi^* or *Prkcd ^f/f^ Cd19^cre/+^ B1-8^hi^* mice were labeled with CTV and transferred into B6 mice, which were immunized with NP-Ficoll on the next day. Spleen cells were analyzed three days later. (A) Representative flow cytometric plots of the spleen cells with the numbers indicating percentages of the cells within the neighboring gates (left). The frequency of IgG3^+^ cells among whole donor B cells (CD45.1^+^ CD19^+^ CD138^-^) (middle) and at each cell division number (right) are shown. (B) qRT-PCR analysis of the indicated transcripts in the donor B cells collected as in Figure 4 - figure supplement 1A. **(C)** *Prkcd^+/+^ Cd19^cre/+^ B1-8^hi^* or *Prkcd ^f/f^ Cd19^cre/+^ B1-8^hi^* B cells transduced with an empty vector (Ev) or vectors expressing AID or PKCδ were transferred into B6 mice that had been immunized with NP-Ficoll on the previous day as in Figure 4 - figure supplement 1B. Representative flow cytometry plots of the donor cells transduced with the vectors (CD45.1^+^GFP^+^, gated as in Figure 4 - figure supplement 1C) (left) and the frequency of IgG3^+^ cells among the CD45.1^+^GFP^+^ cells (right) on day 3 after transfer. Small horizontal bars are the means of 5 (A) or 6-9 (C) biological replicates. Each symbol represents an individual mouse (A, middle; C). The symbols are the means of 5 biological replicates (A, right). Data are means ± SDs of 3 technical replicates pooled from 5 mice (B). The data are representative of 3 (A) or 2 (B) independent experiments or is pooled from 2 independent experiments (C). ns, not significant (p > 0.05); **, p<0.01; ****, p<0.0001; p values were calculated by unpaired Multiple t test (A) or 1-way ANOVA with Tukey’s test (C). The following source data and figure supplement are available for Figure 4: Source data 1. Source data for Figure 4A-C. Figure supplement 1. Supplementary data for Figure 4.

We next analyzed molecular events associated with CSR in sorted donor cells (Figure 4 - figure supplement 1A). The amounts of the Iγ3-Cµ circle transcripts and the Iµ-Cγ3 early post-switch transcripts were far less in *Cd19^cre/+^ Prkcd^f/f^* B cells compared to control cells, indicating fewer CSR events in the former *in vivo* (Figure 4 B). The expression of Iµ-Cµ and Iγ3-Cγ3 germline transcripts were comparable between the *Cd19^cre/+^ Prkcd^f/f^* B cells and the control cells, whereas the expression of *Aicda* was substantially reduced in the former (Figure 4 B). These results indicate that PKCδ mediates CSR to IgG3 through upregulation of AID mRNA, but not through the activation of *Ig* gene S regions in B cells responding to TI-2 Ags *in vivo*.

To examine whether the reduction of AID is a primary reason for the impaired generation of IgG3^+^ cells from PKCδ-deficient B cells, we transduced AID into *in vivo* primed *Cd19^cre/+^ Prkcd^f/f^ B1-8^hi^* B cells and transferred them into B6 mice that had been immunized with NP-Ficoll one day previously (Figure 4 - figure supplement 1B). The reconstitution of AID expression restored IgG3 class switching to a frequency comparable to the same cells reconstituted with PKCδ (Figure 4 - figure supplement 1C and Figure 4 C). Collectively, these results indicate that PKCδ mediates class switching to IgG3 by upregulating the expression of AID in the TI-2 response.

### PKCδ upregulates the transcription of *AID* through BATF

It has been shown that expression of *Aicda* gene is regulated by various transcriptional factors (Vaidyanathan et al., 2014; Tran et al., 2010; Crouch et al., 2007). To assess the role of PKCδ in *Aicda* gene expression, we quantified expression levels of the genes encoding such transcription factors in PKCδ-sufficient and PKCδ-deficient B cells collected from mice immunized with NP-Ficoll. Among the genes tested, we found that the amount of *Batf* mRNA was markedly lower in *Cd19^cre/+^ Prkcd^f/f^* B cells than in control cells (Figure 5 A). The expression of BATF mRNA and protein was induced by *in vitro* stimulation with NP-Ficoll alone in PKCδ-sufficient B cells, but only marginally in PKCδ-deficient B cells (Figure 5, B and C). *Batf* expression was not induced by IL-1α, IL-1β, IFNα, or NP-CGG in either group (Figure 5 B) and BATF protein was not induced by NP-CGG alone (Figure 5 - figure supplement 1A). Collectively, these data indicate that PKCδ induces the expression of BATF downstream of the BCR in the TI-2 response.

**Figure 5.**
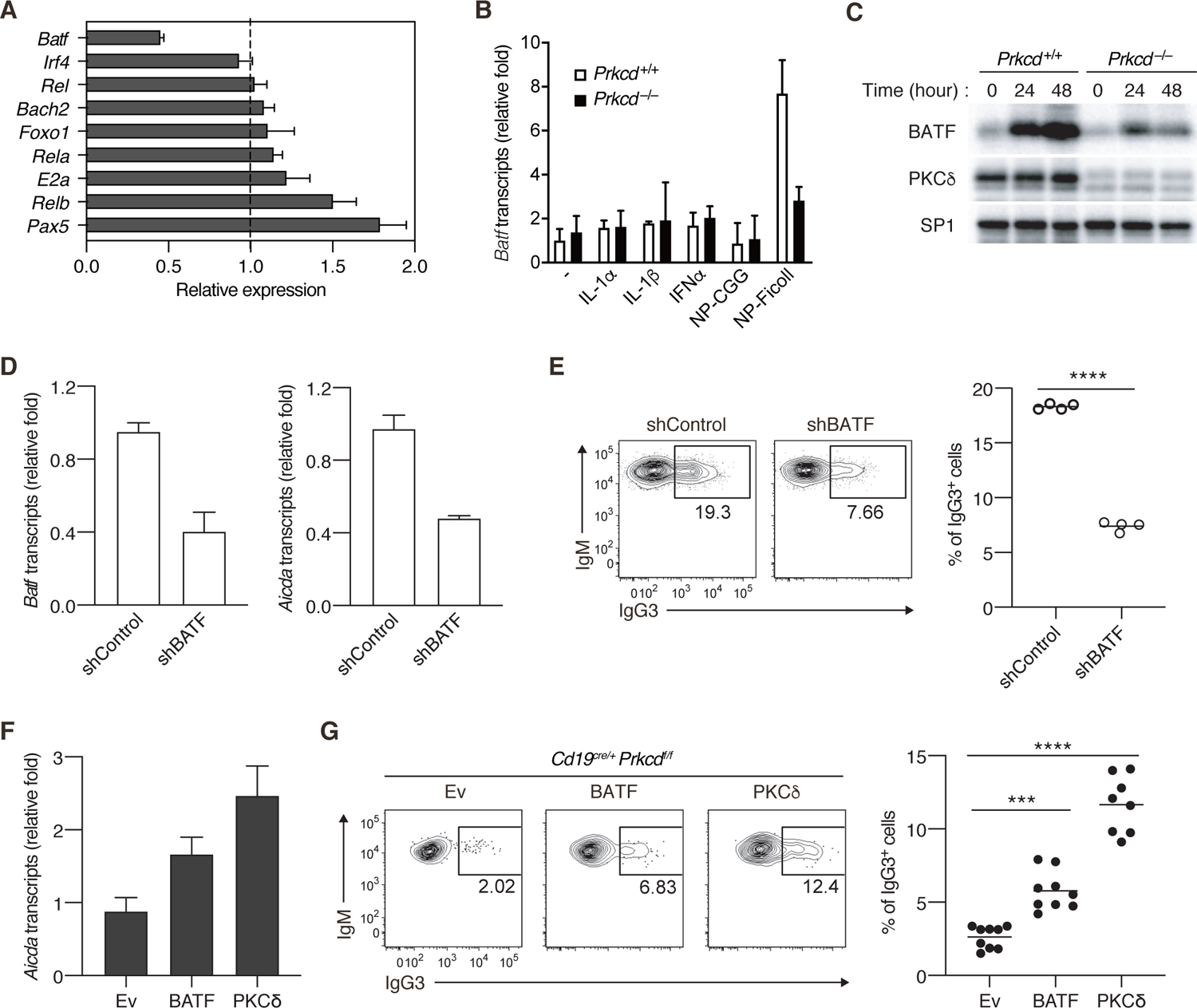
PKCδ regulates IgG class switching by upregulating BATF expression. **(A)** qRT-PCR analysis of the transcripts of the indicated genes in *Prkcd^+/+^ Cd19^cre/+^ B1-8^hi^* or *Prkcd ^f/f^ Cd19^cre/+^ B1-8^hi^* donor B cells (CD45.1^+^CD19^+^CD138^-^) purified as in Figure 4 - figure supplement 1A from the recipient mice immunized with NP-Ficoll 3 days previously. Shown is the relative expression of each gene in *Prkcd ^f/f^ Cd19^cre/+^* cells to that in *Prkcd ^+/+^ Cd19^cre/+^* cells. **(B)** qRT-PCR analysis of *Batf* transcripts in *Prkcd* ^+*/+*^ *Igκ^-/-^ B1-8^flox/+^* or *Prkcd ^-/-^ Igκ^-/-^ B1-8^flox/+^* B cells cultured with medium alone (–) or with the indicated stimuli for 2 days. **(C)** Immunoblot analysis in *Prkcd* ^+*/+*^ *Igκ^-/-^ B1-8^flox/+^* or *Prkcd ^-/-^ Igκ^-/-^ B1-8^flox/+^* B cells stimulated with NP-Ficoll for the indicated times. **(D and E)** *B1-8^hi^* B cells transduced with knockdown vectors for luciferase (shControl) or BATF (shBATF) were transferred into B6 mice that had been immunized with NP-Ficoll on the previous day as in Figure 4 - figure supplement 1B, and their spleen cells were analyzed on day 4 after transfer. (D) qRT-PCR analysis of *Aicda* and *Batf* transcripts in the vector-transduced donor B cells (CD45.1^+^ GFP^+^ CD19^+^ CD138^-^) collected as in Figure 4 - figure supplement 1C. (E) Representative flow cytometric plots of the transduced donor cells with the numbers indicating the percentage of IgG3^+^ cells (left) and the frequency of the IgG3^+^ cells among such cells (right) gated as in Figure 4 - figure supplement 1C. **(F and G)** *Prkcd ^f/f^ Cd19^cre/+^ B1-8^hi^* B cells transduced with Ev or vectors expressing BATF or PKCδ were transferred into B6 mice that had been immunized with NP-Ficoll on the previous day, and their spleen cells were analyzed 3 days after transfer. (F) qRT-PCR analysis of *Aicda* transcripts in the vector-transduced donor B cells. (G) Representative flow cytometric plots of the transduced donor cells with the numbers indicating the percentage of IgG3^+^ cells (left) and the frequency of the IgG3^+^ cells among such cells (right). Data are means ± SDs of 3 technical replicates (A, B, D, F). Samples were pooled from 5-8 mice (D and F). Small horizontal bars are the means of 4 (E) or 8-9 (G) biological replicates. Each symbol represents an individual mouse (E and G). The data are representative of 2 independent experiments (A-F) or is pooled from 2 independent experiments (G). ***, p<0.001; ****, p<0.0001; p values were calculated by 2-tailed unpaired Student’s t test (E) or 1-way ANOVA with Tukey’s test (G). The following source data and figure supplement are available for Figure 5: Source data 1. Source data for Figure 5A, B, D-G; Source data 2-5. Source data for Figure 5C. Figure supplement 1. Supplementary data for Figure 5.

It was reported that BATF binds to a regulatory region of the *Aicda* gene to directly promote its expression and that IgG3 production against TNP-Ficoll was impaired in *Batf^-/-^* mice (Ise et al., 2011). Therefore, we next asked whether the defect of BATF expression is responsible for the suppression of AID expression and IgG3^+^ cell generation in PKCδ-deficient mice. First, we knocked down BATF in *B1-8^hi^* B cells and transferred them into mice immunized with NP-Ficoll. Both the expression of *Aicda* and the frequency of IgG3^+^ cells were significantly decreased in BATF knock-down cells compared with the mock-transduced control cells (Figure 5, D and E). Conversely, forced expression of BATF in *B1-8^hi^ Cd19^cre/+^ Prkcd^f/f^* B cells partially but significantly restored *Aicda* expression and the generation of IgG3^+^ cells in the recipient mice immunized with NP-Ficoll (Figure 5, F and G). Taken together, these data indicate that PKCδ mediates expression of AID and class switching to IgG3 through upregulation of BATF expression in B cells undergoing a TI-2 response.

### PKCδ is required for homeostatic anti-bacterial IgG3 production and protection against bacteremia

Recent works have revealed that commensal microbes induce an IgG response and confer protection against systemic bacterial infection (Zeng et al., 2016). Among anti-bacterial IgG, IgG3 is most abundant and produced in a T cell-independent manner (Ansaldo et al., 2019; Koch et al., 2016). Therefore, we assessed the contribution of PKCδ in IgG-mediated anti-bacterial responses. To standardize the microbiota, we cohoused control and *Cd19^cre/+^ Prkcd^f/f^* mice over four weeks and serum antibodies against fecal bacteria were titrated. Production of serum anti-bacterial IgM, IgG1 and IgG2b was not changed significantly, but that of IgG3 was severely impaired in *Cd19^cre/+^ Prkcd^f/f^* mice (Figure 6 A). Anti-bacterial IgG2c was undetectable in both mouse groups (data not shown). Given that PKCδ was required for production of all IgG subclasses in the anti-NP TI-2 response (Figure 3), the PKCδ-independent production of anti-bacterial IgG1 and IgG2b may be attributable to TD responses, as reported for IgG1 (Ansaldo et al., 2019), while anti-bacterial IgG3 is mainly produced by the TI-2 response.

**Figure 6.**
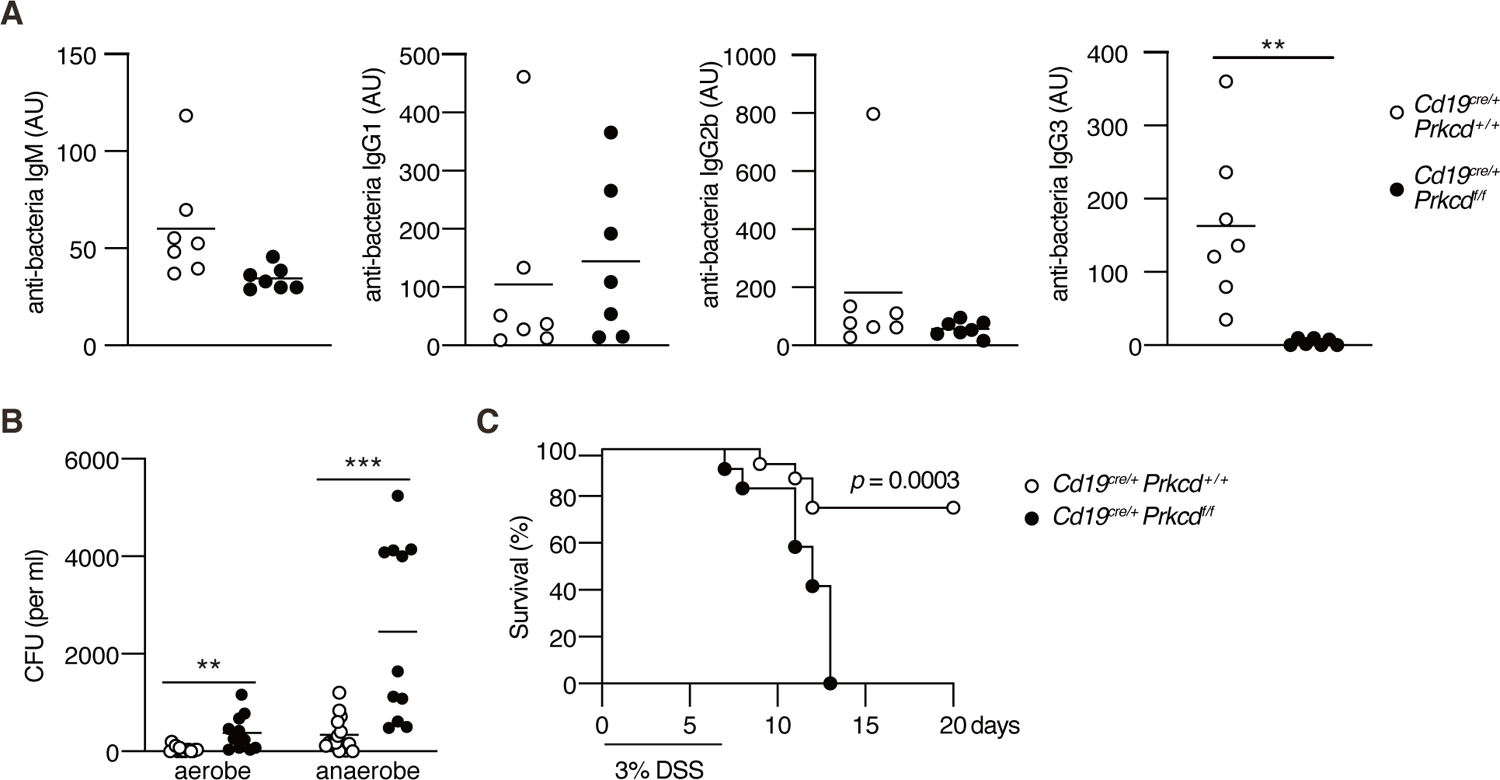
PKCδ is required for protective anti-bacterial IgG3 production *Cd19^cre/+^ Prkcd^+/+^* and *Cd19^cre/+^ Prkcd ^f/f^* mice were co-housed for at least 4 weeks and treated with 3% DSS for 7 days. **(A)** Serum IgM, IgG1, IgG2b, and IgG3 against fecal bacteria after co-housing were quantified by ELISA. AU, arbitrary units. **(B)** Colony forming unit (CFU) of aerobes and anaerobes in the blood 7 days after DSS treatment. **(C)** Kaplan-Meier survival plot of 16 (*Cd19^cre/+^ Prkcd^+/+^*) and 12 (*Cd19^cre/+^ Prkcd^f/f^*) mice at indicated days. Small horizontal bars are the means of 7 (A), or 15 (*Cd19^cre/+^ Prkcd^+/+^*) or 11 (*Cd19^cre/+^ Prkcd^f/f^*) in (B), biological replicates. Data were obtained from 16 (*Cd19^cre/+^ Prkcd^+/+^*) or 12 (*Cd19^cre/+^ Prkcd^f/f^*) mice in (C). Each symbol represents an individual mouse (A and B). The data is representative of 2 independent experiments (A) or pooled from 2 independent experiments (B and C). **, p<0.01; ***, p<0.001; p values were calculated by 2-tailed unpaired Welch’s t test (A), unpaired Multiple t test (B), or Log-rank test (C). The following source data is available for Figure 6: Source data 1. Source data for Figure 6A, B.

We further asked whether regulation of commensal bacteria is defective in the PKCδ-deficient mice. Dextran sodium sulfate (DSS) treatment is known to disrupt the gut epithelium and to allow intestinal bacteria to translocate throughout the body. Subsequently, it leads to fatal bacteremia in the absence of microbiota-specific IgG (Zeng et al., 2016). After the treatment with DSS, *Cd19^cre/+^ Prkcd^f/f^* mice exhibited increased numbers of aerobic and anaerobic bacteria in the blood compared to control mice (Figure 6 B). Accordingly, the mortality of *Cd19^cre/+^ Prkcd^f/f^* mice was significantly higher than that of control mice (Figure 6 C). Collectively, these results suggest that IgG3 production by a TI-2 response via PKCδ prevents lethal bacteremia.

## Discussion

Although previous reports showed that proximal BCR signal molecules such as BLNK or Btk are necessary for B cell activation and subsequent IgM and IgG production in the TI-2 response in mice (Xu et al., 2000; Ellmeier et al., 2000), it was not clear whether there are any specific BCR signaling pathways inducing CSR. Here, we demonstrated that BCR stimulation with a TI-2 Ag (NP-Ficoll), but not a TD Ag (NP-CGG), both sharing the same BCR epitope, promoted transcription of AID and induced IgG3 CSR in the presence of a secondary stimulation. BCR engagement with a TI-2 Ag induced the phosphorylation of PKCδ and PKCδ was required for upregulation of BATF expression, thereby mediated the induction of *Aicda* transcription and subsequent CSR to the IgG subclasses (Figure 7). By contrast, PKCδ was dispensable for TI-2 Ag-mediated B cell proliferation, IgM production, as well as for B-cell development and the TD response. Thus, we have found for the first time, as far as we know, a BCR signaling molecule that selectively mediates induction of CSR in the TI-2 response.

**Figure 7.**
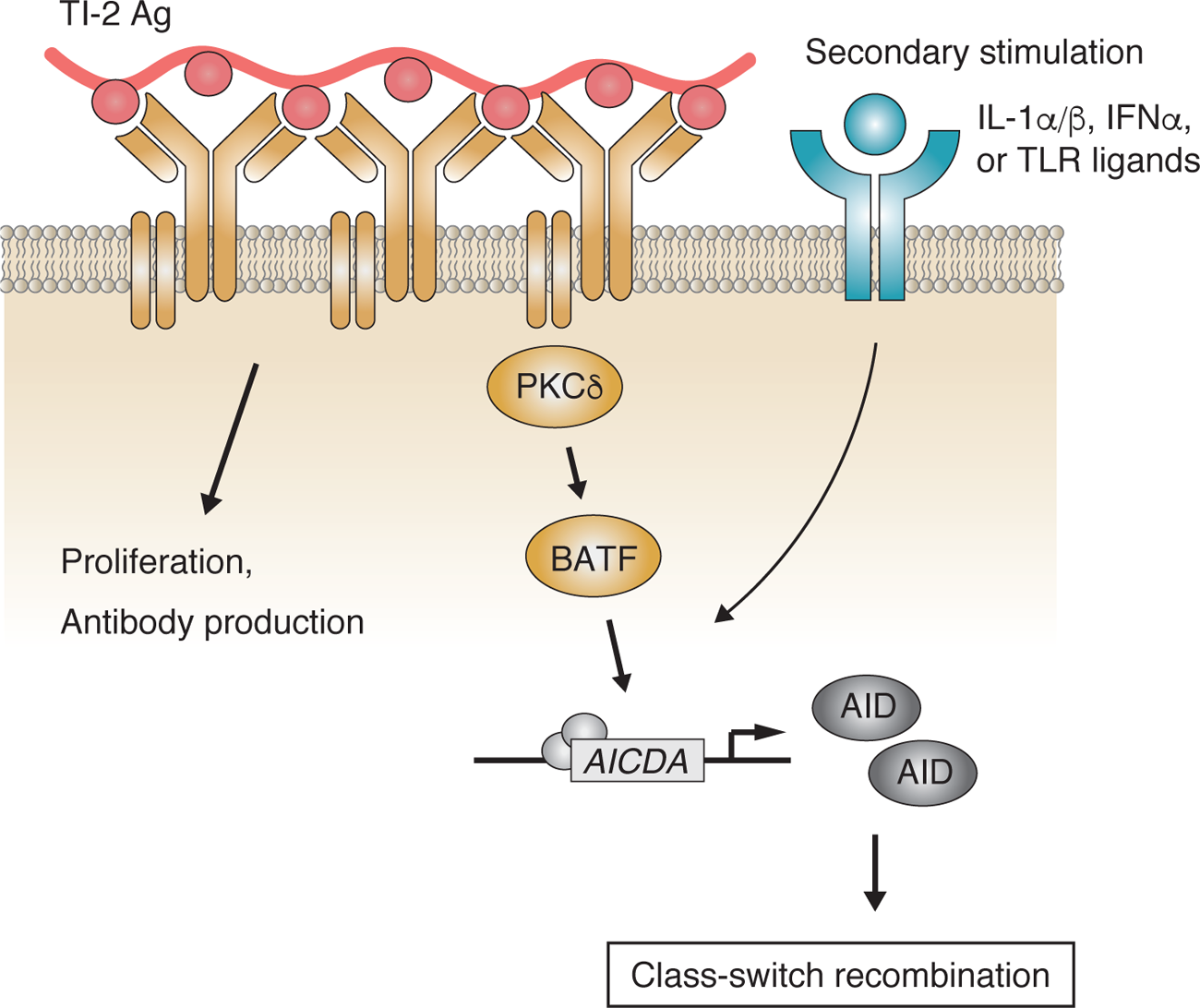
Working model of the PKCδ-mediated induction of CSR in the TI-2 response BCR stimulation with a TI-2 Ag activates PKCδ signaling and induces the expression of BATF, which works cooperatively with secondary signals induced by IL-1α/β, IFNα, or TLR ligands and drives the transcription of AID to induce IgG CSR.

TI-2 Ags such as NP-Ficoll can induce antibody response in the absence of major histocompatibility complex (MHC) class II-restricted T cell help or MyD88 mediated TLR receptor signaling (Gavin et al., 2006; Hou et al., 2011). Thus, it has been unclear whether any co-stimulation is necessary for B cell activation in a TI-2 response. We demonstrated that NP-Ficoll, but not NP-CGG, was able to activate NP-specific naive B cells to induce proliferation, antibody production and potentiation for CSR *in vitro*. Thus, despite sharing the same antigenic epitope (NP) at a similar average number per molecule, these Ags appear to elicit disparate BCR signal transduction. After Ag binding, the BCR is clustered and recruited to lipid microdomains, in which signaling proteins are localized that trigger B cell activation signaling (Niiro and Clark, 2002). Ag structure probably affects such BCR dynamics or membrane organization of signal components. Consistently, it was previously reported that a high-valency TI-2 Ag induces the formation of large BCR clusters in the lipid microdomains (Puffer et al., 2007).

We have revealed that, in the TI-2 response, BCR-downstream signaling through PKCδ is critical for CSR and generation of IgG. PKCδ is known to control Ag-induced tolerance in immature B cells (Mecklenbräuker et al., 2002; Limnander et al., 2011). However, a possible alteration of the mature B-cell repertoire in PKCδ-deficient mice is not attributable to the defective class switching to IgG3, since the defect was obvious in the NP-Ficoll response by PKCδ-deficient, B1-8V_H_-knock-in B cells with a monoclonal anti-NP repertoire. Although PKCδ mediates BCR-induced Erk signaling in immature B cells (Limnander et al., 2011), the phosphorylation of Erk was normal in PKCδ-deficient mature B cells stimulated with NP-Ficoll (data not shown). Accordingly, the function of PKCδ in B cells might be different across developmental stages and the role of PKCδ in BCR signaling in mature B cell responses has been unknown until now.

Here, we demonstrated that PKCδ mediated the expression of AID to induce CSR. BCR engagement with a TI-2 Ag induced the expression of BATF via PKCδ and BATF was required for the transcription of the *Aicda* gene in a TI-2 response, as reported previously (Ise et al., 2011). Forced expression of BATF in PKCδ-deficient B cells restored the expression of AID and IgG3 class switching significantly, but less effectively, than that of PKCδ itself. In this regard, various transcription factors have been reported to cooperatively induce the transcription of AID in the TD response (Vaidyanathan et al., 2014; Crouch et al., 2007; Tran et al., 2010). Although we have not assessed the contribution of other transcription factors in the TI-2 response, since the expression of other candidate genes was intact in PKCδ-deficient B cells (Figure 5 A), PKCδ signaling might regulate other transcription factors in addition to BATF to fully upregulate the expression of AID to induce CSR in the TI-2 response.

On the other hand, PKCδ was not required for IgG production in the TD response, like other proximal BCR molecules (Khan et al., 1995; Xu et al., 2000). *In vitro* stimulation with a TD Ag did not induce the phosphorylation of PKCδ, the expression of BATF, and AID or class switching. Thus, BCR signaling via PKCδ seems to function in CSR only in the TI response.

The synergistic effect of TLR and BCR on the induction of AID has been shown previously (Pone et al., 2012). Here we found that IL-1α/β, IFNα, and TLR ligands co-stimulate B cells with TI-2 Ag to induce class switching to IgG3 and perhaps other IgG subclasses, while other cytokines tested barely induced IgG3 production *in vitro*. Although IL-4 synergizes with CD40L to induce IgG1 CSR in the TD-response (Dedeoglu, 2004), IL-4 did not induce IgG3 or even IgG1 production with NP-Ficoll (Figure 1 - figure supplement 1D, data not shown). Consistent with our *in vitro* data, previous reports showed the contribution of IL-1α/β, IFNα, and TLR ligands in several types of TI-2 responses: IgG production upon immunization with NP-Ficoll was dampened in IL-1α/β double-deficient mice (Nakae et al., 2001), and that with NP-Ficoll plus poly:IC was dependent on the IFNα receptor on B cells (Swanson et al., 2010). While IgG production against NP-Ficoll did not require MyD88-mediated TLR signaling (Gavin et al., 2006; Hou et al., 2011), IgG production against pneumococcal polysaccharides requires TLR signaling (Sen et al., 2005). Compared to NP-Ficoll, physiological TI-2 Ags are more complex (Snapper, 2006) and may contain pathogen-associated molecular patterns (PAMPs) that mediate TLR signaling. Therefore, any of IL-1α/β, IFNα, and TLR ligands could function as co-stimulation for an *in vivo* TI-2 response, depending on the type of Ag.

Our data revealed that PKCδ is generally required for IgG production in response to TI-2 Ags such as NP-Ficoll and PPS3. Given the predominant role of IgG3 in anti-pneumococcal responses (McLay et al., 2002), PKCδ seems to be needed for protection from pneumococcal infection. Furthermore, we found that PKCδ is required to produce IgG3 against commensal bacteria in the steady-state and the prevention of bacteremia after epithelial barrier disruption. As commensal bacteria have been shown to contain various antigens and stimulate multiple immune pathways (Belkaid and Harrison, 2017), our result suggests that some commensal bacteria express TI-2 Ag-like repetitive structures that elicit IgG3 production. It has been shown that such IgG against symbiotic bacteria also plays a protective role in systemic infection by pathogens (Zeng et al., 2016). Taken together, IgG production in the TI-2 response seems to be critical for regulation of various types of bacterial invasion. Loss-of-function PKCδ mutations in humans cause SLE-like autoimmunity, likely due to abnormal B cell tolerance mechanisms (Salzer et al., 2016). Some of these patients also show reduced IgG positive B cells in the peripheral blood and have recurrent infections (Kiykim et al., 2015; Salzer et al., 2013; Kuehn et al., 2013), the reason for which remains unknown. Thus, PKCδ-mediated IgG production by the TI-2 response also appears to be critical in humans for host defense against certain bacteria.

## Materials and methods

### Mice and immunizations

C57BL/6NCrSlc (B6) mice were purchased from Japan SLC. All the following mice were backcrossed to the B6 or B6-CD45.1 strain: *B1-8^flox/+^* mice (Lam et al., 1997), *B1-8^hi^* mice (Shih et al., 2002), *Igκ^-/-^* mice (Chen et al., 1993), *Cd19^cre/+^* mice (Rickert et al., 1995), *Prkcd^-/-^* mice (Miyamoto et al., 2002). *Prkcd ^fl/fl^* mice on the B6 background were developed by Drs. Niino, Shioda and Sakimura (Li et al., 2020) and purchased from the RIKEN BioResource Center (RBRC06462). Mice were immunized i.p. with 100 µg of NP_46_-Ficoll (F-1420; Biosearch Technologies), 1µg of PPS3 (169-X; American Type Culture Collection) or 100 µg of NP_40_-CGG in alum (Haniuda et al., 2016) unless otherwise noted. For flow cytometry of spleen cells, mice were immunized i.v. with 100 µg of NP_46_-Ficoll. Sex-matched 7-14 week-old mice were used for all experiments. All mice were maintained in the Tokyo University of Science (TUS) mouse facility under specific pathogen-free conditions. Mouse procedures were performed under protocols approved by the TUS Animal Care and Use Committee.

### B-cell culture

Spleen cells were stained with a cocktail of biotinylated Abs for CD4, CD8, CD43, CD49b, Ter119 and Streptavidin Particles Plus DM, from which naive B cells were purified by magnetic negative sorting using the IMag system (BD Biosciences) and MACS system (Miltenyi Biotec), as described previously (Nojima et al., 2011). B cells were cultured in RPMI-1640 medium (Wako) supplemented with 10% heat-inactivated fetal bovine serum (FBS), 10 mM HEPES pH 7.5, 1 mM sodium pyruvate, 50 mM 2-mercaptoethanol, 100 U/ml penicillin, and 100 mg/ml streptomycin (GIBCO). Typically, B cells were cultured at 2×10^5^/ml in the presence of the following stimuli at the indicated doses, unless otherwise noted: NP_46_-Ficoll (10 ng/ml), NP_40_-CGG (10 ng/ml), LPS (1 µg/ml, L2880; Sigma), R-848 (1 µg/ml, tlrl-r848; InvivoGen), CpG ODN 1826 (1 µg/ml, tlrl-1826; InvivoGen), IL-1α (1 ng/ml, 211-11A; Pepro Tech), IL-1β (1 ng/ml, 211-11B; Pepro Tech) or IFNα (100 ng/ml, 752802; Biolegend). Cytokines used in Figure 1 - figure supplement 1D were added at the concertation shown in Supplementary Table 1.

### Retroviral Transduction

To produce retrovirus, pSIREN- or pMXs-based plasmids were co-transfected together with pVSVG into Plat-E cells (kindly provided by T. Kitamura, University of Tokyo) using PEI Max (Mw 40,000, 24765-1; Polysciences). The virus-containing supernatant was harvested 2 days after transfection. For retroviral transduction, B cells were pre-activated *in vivo*: *B1-8^hi^* mice were injected i.p. with 50 µg of NP-Ficoll, and then B cells were purified from the spleens of these mice on the next day. These B cells were mixed with the virus-containing supernatant and spin-infected at 2,000 rpm, 37°C for 90 min with 10 mg/ml DOTAP Liposomal Transfection Reagent (11202375001; Sigma). One day later, the cells were harvested and 5×10^5^ cells were transferred into B6 mice that had been immunized i.v. with NP-Ficoll on the previous day. This strategy is summarized in Figure 4 - figure supplement 1B.

### Proliferation assay

The proliferation assay was performed as described previously (Fukao et al., 2014). Naive B cells were cultured at 5×10^4^ cells/well in 96-well plates for 72 hours, with the last 8 hours in the presence of [^3^H] thymidine (1 mCi/well, NET027001MC; PerkinElmer). Incorporated [^3^H] thymidine was counted by a BetaPlate scintillation counter (Wallac, Gaithersburg, MD).

### Flow cytometry

Single-cell suspensions from spleen or peritoneal cavity were prepared, red blood cells were lysed with ammonium chloride buffer and then cells were incubated with anti-CD16/32 Ab (2.4G2) to block FcγRs. Cultured B cells were collected in MACS buffer (PBS supplemented with 0.5% BSA, 2 mM EDTA) at the indicated days of culture. Cells were stained with Abs and reagents on ice (for splenocytes) or at room temperature (for cultured B cells). For the staining of IgM, IgG, and NP-binding Ig, Fixation/Permeabilization Solution Kit (554714; BD Biosciences) was used according to the manufacturer’s protocol to detect total (surface and intracellular) proteins, after surface staining of other molecules. NP-binding Ig was stained with NP_14_-BSA-Alexa Fluor 647 (Haniuda et al., 2016). Dead cells were stained with Fixable Viability dye eFluor 506 (65-0866-18; Thermo Fisher Scientific) before cell fixation and excluded from analysis. All samples were analyzed using FACSCanto II, FACSAria II or III (BD Biosciences) with FlowJo software (Tree Star, Inc.). Antibodies used in this study are listed in Supplementary Table 2.

### Cell division analysis

Naïve B cells were resuspended in PBS at 5×10^6^ cells/ml and labeled with 5 µM of CellTrace^TM^ Violet (CTV) Cell proliferation Kit (C34557; Thermo Fisher Scientific) at 37°C for 20 minutes according to the manufacturer’s protocol. Collected cells were analyzed by flow cytometry as described above. Cell divisions were determined using the proliferation platform of FlowJo.

### Adaptive transfer and donor B cell purification

1×10^6^ CD45.1 *B1-8^hi^* naïve B cells were transferred into B6 mice, which were then immunized i.v. with NP-Ficoll on the next day. Donor B cells were magnetically enriched from pooled spleens of recipient mice using APC-conjugated anti-CD45.1 and anti-APC MicroBeads (130-090-855; Miltenyi Biotec), with the MACS system (Miltenyi Biotec). After enrichment, cells were further stained with respective Abs and sorted using FACSAria II or III (BD Biosciences) as shown in Figure 4 - figure supplement 1A. Rv-transduced donor B cells were enriched as described above and sorted as shown in Figure 4 - figure supplement 1C.

### RT-PCR, qPCR

TRI Reagent (T9424; Sigma) or RNeasy Micro (74004; QIAGEN) was used to isolate total RNA from B cells. cDNA was generated from total RNA using ReverTra Ace (TRT-101; TOYOBO) with an oligo(dT)20 primer (18418020; Thermo Fisher Scientific) according to the manufacturer’s protocols. For the analysis of a small number of cells, cDNA was generated from cell lysates with SuperPrep II Cell lysis & RT Kit for qPCR (SCQ-401; TOYOBO) according to the manufacturer’s protocols. Quantitative real-time PCR (qPCR) was performed using Thunderbird SYBR qPCR Mix (QPS-201; TOYOBO) with the 7500 fast Real-time PCR system or Quant-Studio 3 (Applied Biosystems). For quantification of gene expression levels, each sample was normalized to the expression of a control housekeeping gene, *Gapdh* or *Rps18*. The relative fold change in expression of each gene compared to a control sample, set as 1.0, was calculated with the 2-ddCT method. Primers used in this study are listed in Supplementary Table 3. The germline and post-switched transcripts were analyzed with the following primer sets: germline Iµ-Cµ transcripts: Iµ Fw1 and Cµ Rv; Iγ3-Cγ3 transcripts: inner Iγ3 Fw and Cγ3 Rv; post-switched Iµ-Cγ3 transcripts: Iµ Fw2 and Cγ3 Rv. For the measurement of the circle Iγ3-Cµ transcript, cDNA was generated with external Cµ Rv primer from total RNA as described above, and pre-amplified with external Iγ3 Fw primer and Cµ Rv primer using GoTaq Green Master Mix (M712B; Promega). Pre-application products were purified with QIAquick PCR Purification Kit (28104; QIAGEN) and circle Iγ3-Cµ transcript was quantified with inner Iγ3 Fw primer and Cµ Rv primer by qPCR as described above. Specific amplification of the circle Iγ3-Cµ transcript was confirmed by analyzing the sequence of the qPCR products in advance. The expression of the Iγ3-Cµ transcript was normalized to the expression of Rps18 in cDNA generated with oligo(dT)20 primer from the same RNA sample.

### ELISA

Concentrations of total IgM, IgG or IgG subclasses and of Ag-specific Igs (where indicated) were assessed by titration of culture supernatants or sera by ELISA. Total and NP-specific antibodies were measured as described previously (Fukao et al., 2014; Nojima et al., 2011), with NP_13.6_-BSA used for coating plates for the latter. PPS3-specific IgM and IgG3 were detected using 96-well plates coated with PPS3. Bacteria-specific antibody was detected as described previously (Zeng et al., 2016). Heat-killed fecal bacteria isolated from *Prkcd^+/+^ Cd19^cre/+^*mice and *Prkcd ^f/f^ Cd19^cre/+^*mice co-housed at least for 4 weeks were mixed and used for plate coating.

### Immunoblotting

Cells were lysed with 1% NP-40 lysis buffer or RIPA buffer (40 mM Tris-HCl pH 7.5, 150 mM NaCl, 1% NP-40, 1% sodium deoxycholate, 0.1% SDS and 1mM EDTA) supplemented with protease and phosphatase inhibitors. Lysates were sonicated and mixed with sample buffer and dithiothreitol (DTT) and boiled. Lysates were resolved on SDS-PAGE and transferred to PVDF membranes (Millipore), followed by immunoblotting as previously described (Haniuda et al., 2020).

### DSS-Induced Bacteremia

*Prkcd^+/+^ Cd19^cre/+^*mice and *Prkcd ^f/f^ Cd19^cre/+^* female mice were co-housed at 1:1 ratio from 4 weeks of age. After at least 4 weeks, co-housed mice were administered 3% DSS (9011-18-1; MP Biomedicals) in drinking water for 7 days and then switched to regular water. On the day of the last DSS administration, blood samples were collected from mice and cultured on Brain Heart Infusion Agar (221570; BD Bioscience) under aerobic or anaerobic conditions for 24 hours to measure CFU.

## Statistical analysis

Biological replication is derived from multiple biological samples (mouse or cell). Technical replication consisted of multiple samples derived from one biological sample. All statistical analyses were performed using GraphPad Prism 8 software. Comparisons between two groups were performed by a two-tailed unpaired Student t test, Welch’s t test (in case F-test is significant: P<0.05), or Multiple t test (for Grouped data). Comparisons between multiple groups were performed by one or two-way ANOVA with Tukey’s multiple comparison. Survival of DSS-treated mice was analyzed by Kaplan-Meier survival plot using Log-rank test. In all cases, *, p < 0.05; **, p < 0.01; ***, p < 0.001; ****, p < 0.0001; and ns, not significant (p > 0.05).

## Acknowledgement

We thank Drs. S. Shioda, Y. Niino, T. Nakamachi, and J. Watanabe (Showa University) for *Prkcd ^fl/fl^* mice; K. Nakayama (Kyushu University) for *Prkcd^-/-^* mice; K. Rajewsky (Max Delbrück Center for Molecular Medicine) for *B1-8^flox/+^* mice and *Cd19^cre/+^* mice; M. Nussenzweig (Rockefeller University) for *B1-8^hi^* mice; F. Alt (Harvard Medical School) and T. Tsubata (Tokyo Medical and Dental University) for *Igκ^-/-^* mice; and P. Burrows for critical reading of the manuscript. This work was supported by the Japan Society for the Promotion of Science (JSPS) KAKENHI (19K16700). The authors declare no competing financial interests. Author contributions: S. Fukao conceptualized the study, designed and performed most of the experiments, analyzed data, and wrote the manuscript. K. Haniuda performed some experiments, analyzed data, edited the manuscript, and gave critical advice. H. Tamaki provided technical support. D. Kitamura supervised the study and wrote the manuscript.

## Additional files

### Supplementary files

Supplementary File 1: Supplementary Tables 1 – 3. The list of cytokines, antibodies and primers used in this study.

## Data availability

All data generated or analyzed during this study are included in the manuscript. Source data are provided for Figures 1-6 and Figure supplements for Figures 1-3, 5 in the Source data files.

## Supplemental Figures

**Figure 1 - figure supplement 1.**
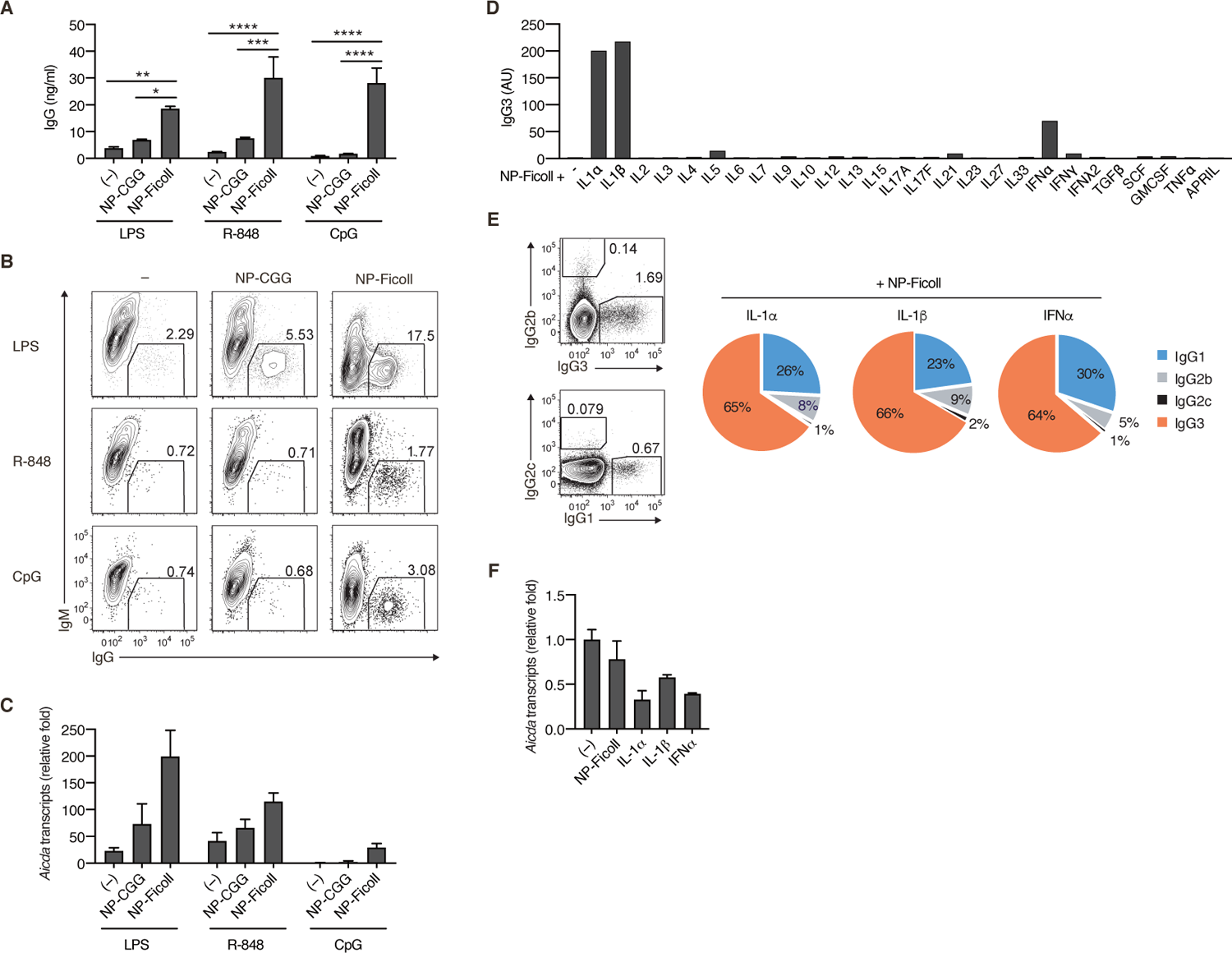
TI-2 Ag induces the generation of IgG^+^ cells in the presence of secondary stimulation. **(A-C)** *Igκ^-/-^ B1-8^flox/+^* naïve B cells were stimulated with nothing (-), NP-CGG or NP-Ficoll in the presence of LPS, R-848 or CpG. (A) IgG concentration in the culture supernatants on day 3. (B) Representative flow cytometric plots on day 3 showing percentages of IgG^+^ B cells. (C) qRT-PCR analysis of *Aicda* transcripts on day 2. **(D and E)** *Igκ^-/-^ B1-8^flox/+^* naïve B cells were stimulated with NP-Ficoll and each indicated cytokine. (D) IgG3 in the culture supernatants on day 4 titrated by ELISA. AU, arbitrary units. (E) Representative flow cytometric profiles of IgG1, IgG2b, IgG2c and IgG3 on day 3 (left) and the ratio of B cells expressing these IgG subclasses estimated from the flow cytometry data (right). **(F)** *Igκ^-/-^ B1-8^flox/+^* naïve B cells were stimulated with nothing (-), NP-Ficoll or the indicated cytokine alone. qRT-PCR analysis of *Aicda* transcripts on day 2. Data are means of 3 biological replicates (E), means ± SDs of 2 biological replicates (A) or 3 technical replicates (C and F). The data are representative of 2 independent experiments. *, p<0.05; **, p<0.01; ***, p<0.001; ****, p<0.0001; p values were calculated by 2-way ANOVA with Tukey’s test (A). The following source data are available for Figure 1 - figure supplement 1: Source data 1. Source data for figure supplement 1A, C, D, F.

**Figure 2 - figure supplement 1.**
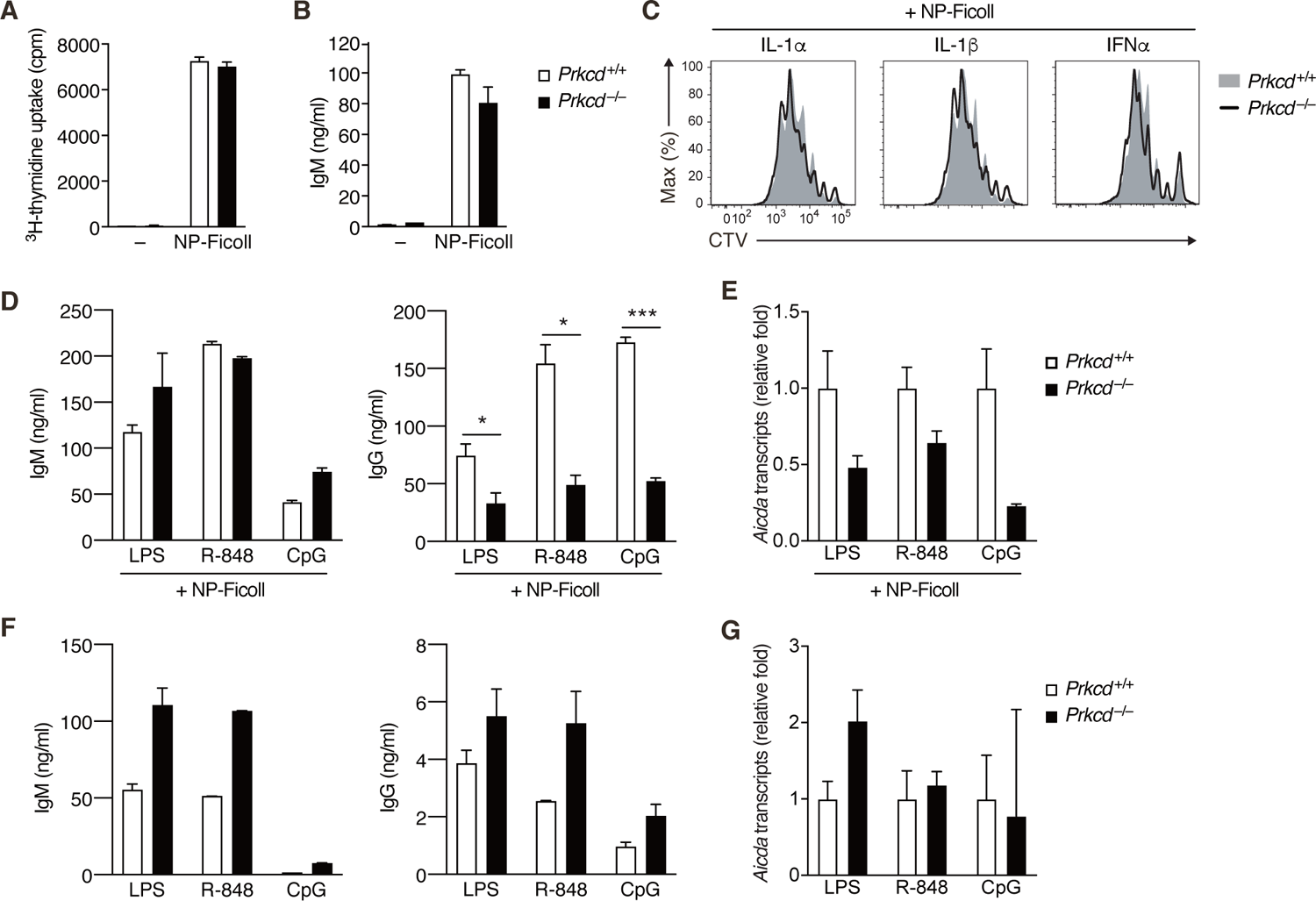
*In vitro* response of PKCδ-deficient B cells. **(A and B)** *Prkcd* ^+*/+*^ *Igκ^-/-^ B1-8^flox/+^* or *Prkcd* ^-/-^ *Igκ^-/-^ B1-8^flox/+^* naïve B cells were cultured with medium alone (–) or NP-Ficoll. (A) ^3^H-thymidine incorporation on day 3. (B) IgM concentration in the culture supernatant on day 4. **(C)** The histogram of CTV in cells analyzed in Fig. 2 C. **(D-G)** *Prkcd*^+*/+*^ *Igκ^-/-^ B1-8^flox/+^* or *Prkcd* ^-/-^ *Igκ^-/-^ B1-8^flox/+^* naïve B cells were cultured with NP-Ficoll and LPS, R- 848 or CpG (D and E) or LPS, R-848 or CpG alone (F and G). (D and F) IgM and IgG concentrations in the culture supernatants on day 3. (E and G) qRT-PCR analysis of *Aicda* transcripts on day 2. Data are means ± SD of 3 (A) or 2-3 (B), 2 (D, F) biological replicates, or 3 technical replicates (E and G). The data are representative of 2 independent experiments. *, p<0.05; ***, p<0.001; p values were calculated by unpaired Multiple t test (A, B, D). The following source data are available for Figure 2 - figure supplement 1: Source data 1. Source data for figure supplement 1A, B, D-G.

**Figure 3 - figure supplement 1.**
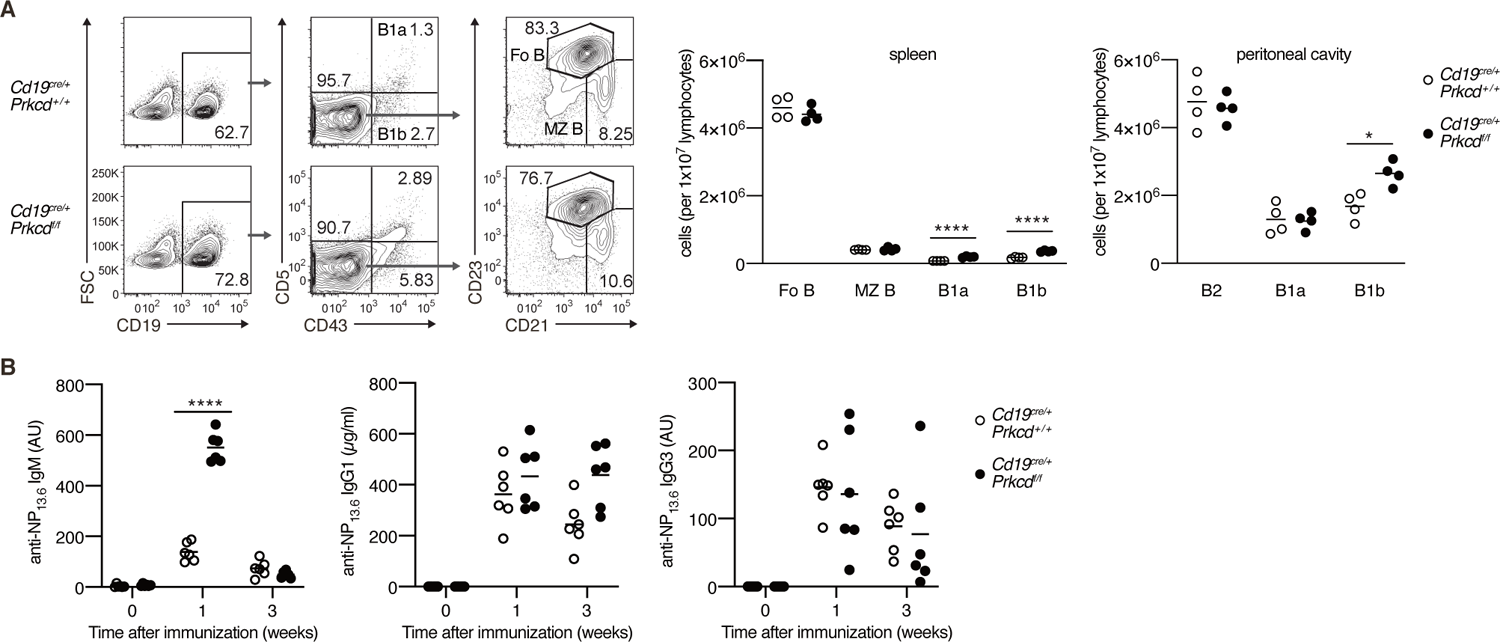
B cell development and TD response of PKCδ-deficient mice. **(A)** Flow cytometric analysis of spleen cells and peritoneal cavity cells of *Cd19^cre/+^ Prkcd^+/+^* and *Cd19^cre/+^ Prkcd ^fl/fl^* mice. Gating strategy of follicular B cells (Fo B; CD19^+^CD43^-^CD21^+^CD23^+^), marginal zone B cells (MZ B; CD19^+^CD43^-^CD21^high^CD23^low^), B1a cells (CD19^+^CD43^+^CD5^+^), and B1b cells (CD19^+^CD43^+^CD5^-^) in the spleen is shown (left). The numbers of Fo B, MZ B, B1a and B1b cells in spleen (middle) and peritoneal cavity (right) are plotted. **(B)** ELISA of anti-NP IgM, IgG1 and IgG3 in the sera of *Cd19^cre/+^ Prkcd^+/+^* and *Cd19^cre/+^ Prkcd ^fl/fl^* mice at indicated weeks after immunization with NP-CGG in alum. AU, arbitrary units. Small horizontal bars are the means of 4 (A) or 6 (B) biological replicates. Each symbol represents an individual mouse. The data are representative of 3 (A and C) or 2 (B and D) independent experiments. *, p<0.05; ****, p<0.0001; p values were calculated by unpaired Multiple t test. The following source data are available for Figure 3 - figure supplement 1: Source data 1. Source data for figure supplement 1A, B.

**Figure 4 - figure supplement 1.**
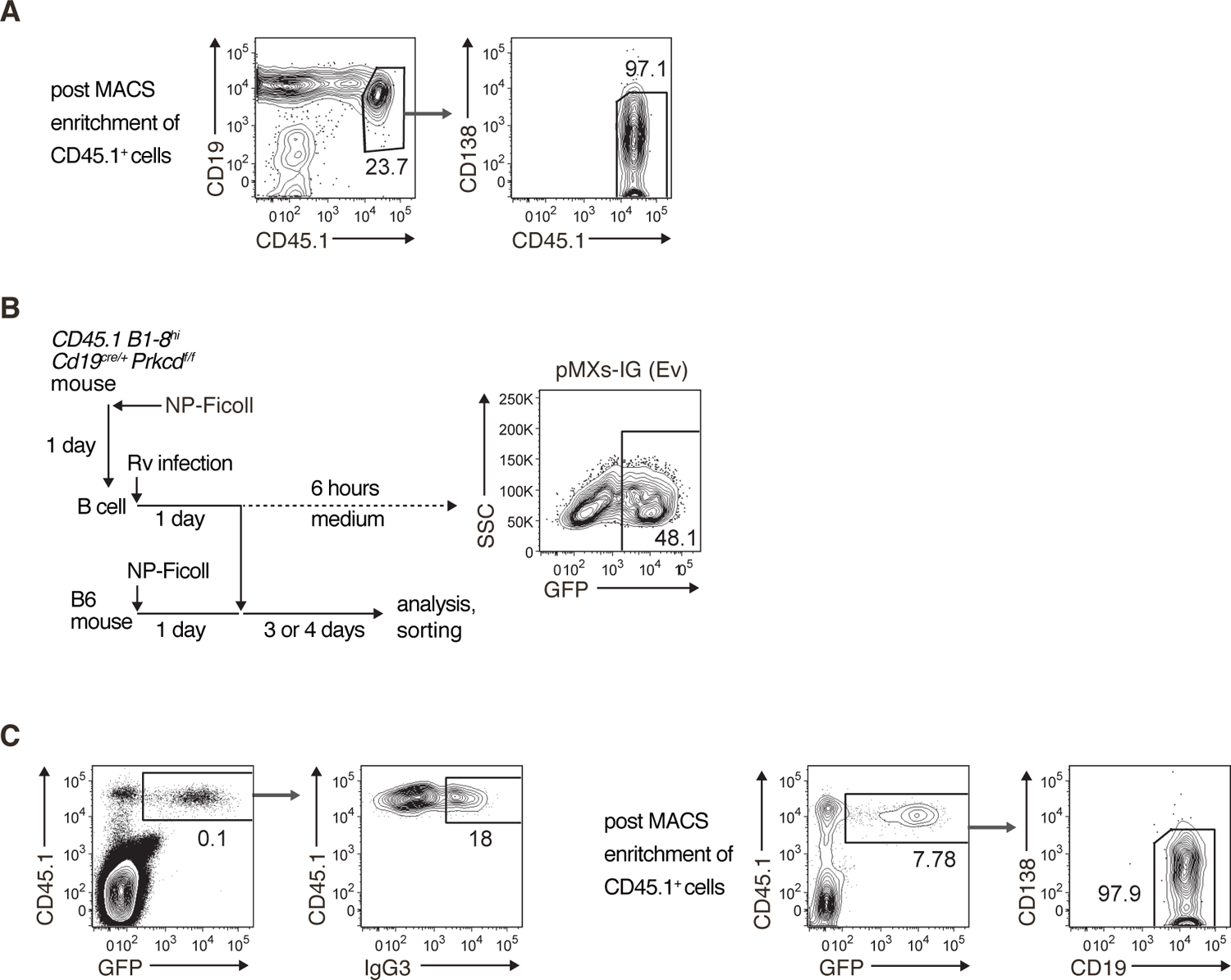
Gating strategies. **(A)** Sorting strategy for donor B cells used in Figure 4 B. The starting population was spleen cells of B6 mice transferred with *Prkcd^+/+^ Cd19^cre/+^ B1-8^hi^* or *Prkcd ^f/f^ Cd19^cre/+^ B1-8^hi^*, CD45.1^+^ B cells and immunized with NP-Ficoll 3 days previously. CD45.1^+^ cells were enriched by a MACS system (left) and then donor B cells (CD45.1^+^CD19^+^CD138^-^) were further sorted by flow cytometry (right). **(B and C)** Strategy for retrovirus (Rv) transduction and the analysis of the TI-2 response (B, left). B cells collected from *Prkcd^+/+^ Cd19^cre/+^ B1-8^hi^* or *Prkcd ^f/f^ Cd19^cre/+^ B1-8^hi^* mice immunized with NP-Ficoll one day previously were infected with Rv and cultured for 1 day. Some Rv-transduced cells were further cultured in medium for 6 hours and analyzed by flow cytometry. Representative data of the cells transduced with pMXs-IRES-GFP empty vector (Ev) is shown (B, right). Other Rv-transduced cells were transferred into B6 mice that had been immunized with NP-Ficoll on the previous day. Spleen cells of recipient mice were analyzed or used for cell sorting 3 or 4 days later. Gating strategy of donor cells transduced with vectors (CD45.1^+^ GFP^+^) and the expression profile of IgG3 are shown (C, left). After the enrichment of CD45.1^+^ cells by a MACS system, the vector-transduced donor B cells (CD45.1^+^ GFP^+^ CD19^+^ CD138^-^) were sorted as shown (C, right). The data are representative of at least 3 independent experiments.

**Figure 5 - figure supplement 1.**
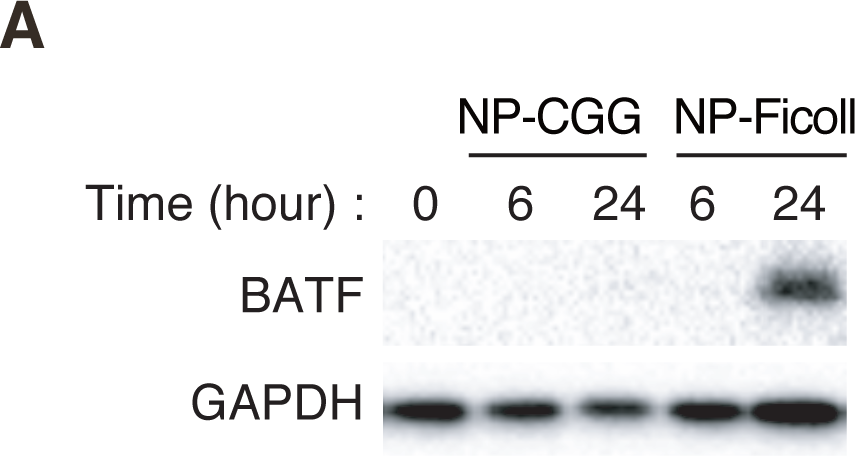
The expression of BATF. **(A)** Immunoblot analysis of BATF and GAPDH in *Igκ^-/-^ B1-8^flox/+^* B cells stimulated with NP-CGG or NP-Ficoll for the indicated times. The data are representative of 2 independent experiments. The following source data are available for Figure 5 - figure supplement 1: Source data 1-3. Source data for figure supplement 1A.

**Table S1.**
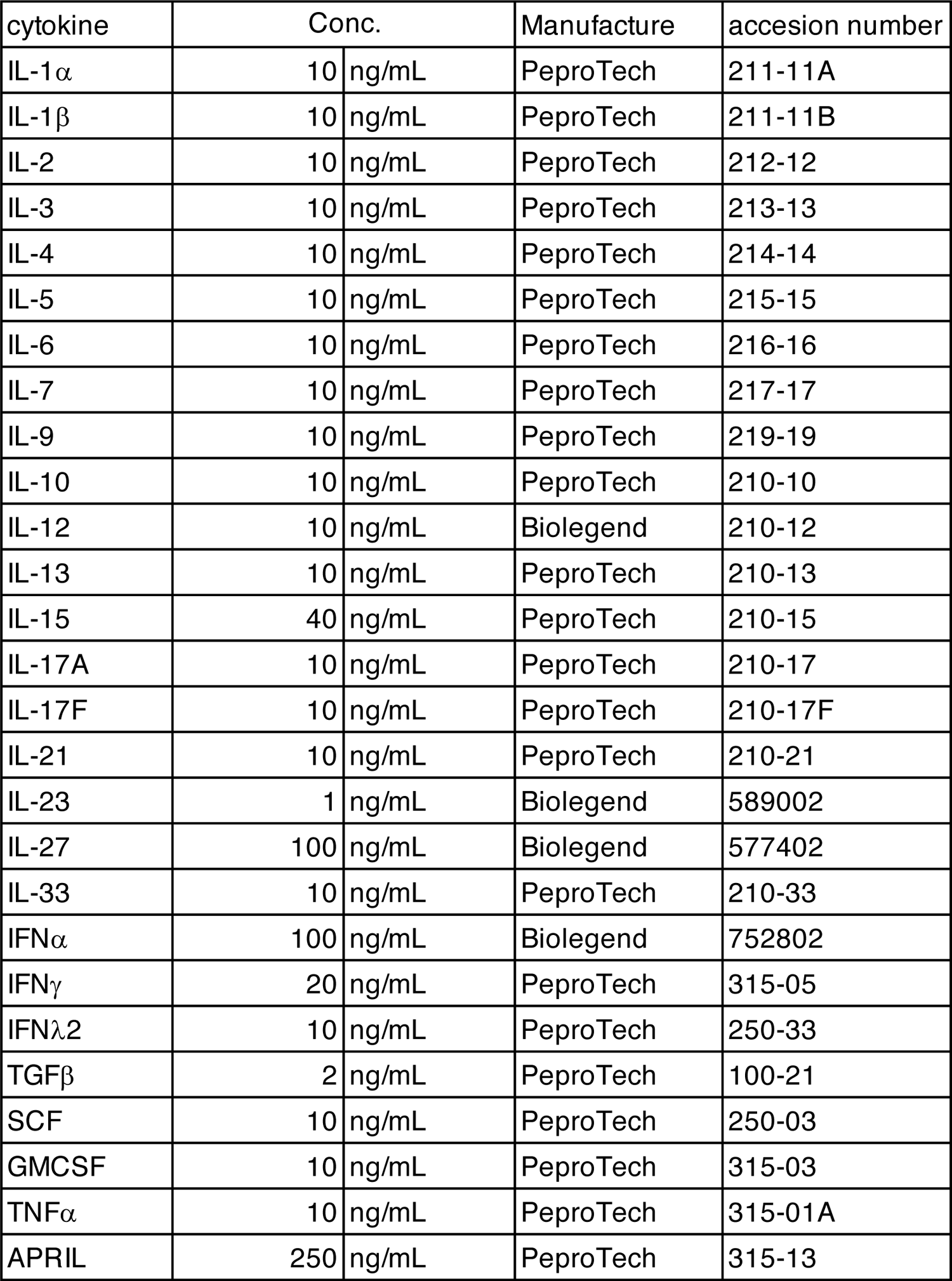
List of cytokines used in Figure S1D

**Table S2.** **List of antibodies used in this study**

| Product name | Manufacture | accession number | Application |
| --- | --- | --- | --- |
| APC-Cy7 anti-mouse CD19 | BioLegend | 115530 | Flow cytometry |
| Biotin Rat Anti-Mouse CD43 | BD Biosciences | 20737 |  |
| BV421 anti-mouse CD138 | BioLegend | 142508 |  |
| BV421 Rat Anti-Mouse IgG3 | BD Biosciences | 565808 |  |
| CD45.1 Monoclonal Antibody (A20), APC | Invitrogen | 17-0453-82 |  |
| FITC anti-mouse CD21/CD35 (CR2/CR1) Antibody | BioLegend | 123407 |  |
| FITC anti-mouse CD5 Antibody | BioLegend | 100605 |  |
| Goat Anti-Mouse IgG2c, Human ads-FITC | SouthernBiotech | 1079-02 |  |
| Goat F(ab')2 Anti-Mouse IgG2b, Human ads-FITC | SouthernBiotech | 1092-02 |  |
| Goat F(ab')2 Anti-Mouse IgG3, Human ads-FITC | SouthernBiotech | 1102-02 |  |
| PE anti-mouse CD23 Antibody | BioLegend | 101607 |  |
| PE anti-mouse IgG2b | BioLegend | 406707 |  |
| PE-Cy7 anti-mouse CD19 | BioLegend | 115520 |  |
| Goat Anti-Mouse IgG Fc-HRP | SouthernBiotech | 1033-05 | ELISA |
| Goat Anti-Mouse IgG, Human ads-UNLB | SouthernBiotech | 1030-01 |  |
| Goat Anti-Mouse IgG1, Human ads-HRP | SouthernBiotech | 1070-05 |  |
| Goat Anti-Mouse IgG1, Human ads-UNLB | SouthernBiotech | 1070-01 |  |
| Goat Anti-Mouse IgG2b, Human ads-HRP | SouthernBiotech | 1090-05 |  |
| Goat Anti-Mouse IgG2b, Human ads-UNLB | SouthernBiotech | 1090-01 |  |
| Goat Anti-Mouse IgG2c, Human ads-HRP | SouthernBiotech | 1079-05 |  |
| Goat Anti-Mouse IgG3, Human ads-UNLB | SouthernBiotech | 1100-01 |  |
| Goat Anti-Mouse IgG3, Human/Bovine/Horse SP ads-HRP | SouthernBiotech | 1103-05 |  |
| Goat Anti-Mouse IgM, Human ads-HRP | SouthernBiotech | 1020-05 |  |
| Goat Anti-Mouse IgM, Human ads-UNLB | SouthernBiotech | 1020-01 |  |
| Anti-GAPDH mAb | MBL | M171-31 | Immunoblotting |
| Anti-SP1 antibody | Abcam | ab227383 |  |
| BATF (D7C5) Rabbit mAb | Cell Signaling Technology | 8638 |  |
| Phospho-PKCdelta (Tyr311) Antibody | Cell Signaling Technology | 2055 |  |
| PKC $\delta$ Antibody (C-17) | Santa Cruz Biotechnology | sc-213 | |

**Table S3.**
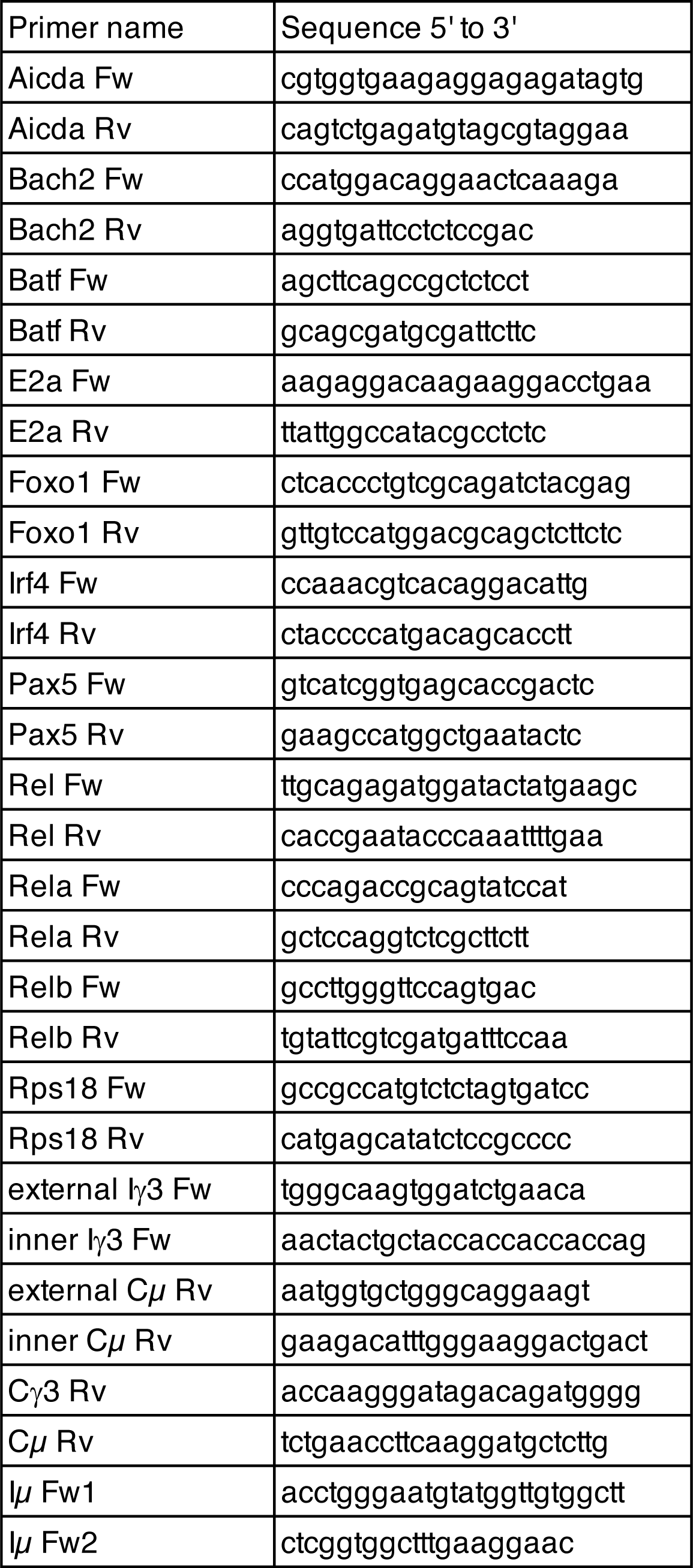
List of primers used in this study

